# Corticohippocampal circuit dysfunction in a mouse model of Dravet syndrome

**DOI:** 10.1101/2021.05.01.442271

**Authors:** Joanna Mattis, Ala Somarowthu, Kevin M. Goff, Jina Yom, Nathaniel P. Sotuyo, Laura M. McGarry, Huijie Feng, Keisuke Kaneko, Ethan M. Goldberg

**Affiliations:** Department of Neurology, The Perelman School of Medicine at The University of Pennsylvania, Philadelphia, PA 19104, USA; Division of Neurology, Department of Pediatrics, The Children’s Hospital of Philadelphia, Philadelphia, PA, 19104 USA; Neuroscience Graduate Group, The University of Pennsylvania Perelman School of Medicine, Philadelphia, PA, 19104 USA; College of Arts and Sciences, The University of Pennsylvania, Philadelphia, PA, 19104 USA; Department of Neuroscience, The Perelman School of Medicine at The University of Pennsylvania, Philadelphia, PA 19104, USA

## Abstract

Dravet syndrome (DS) is a neurodevelopmental disorder defined by treatment-resistant epilepsy, autism spectrum disorder, and sudden death, due to pathogenic variants in *SCN1A* encoding the Nav1.1 sodium channel subunit. Convergent data suggest hippocampal dentate gyrus (DG) pathology. We found that optogenetic stimulation of entorhinal cortex was ictogenic in DS (*Scn1a*^*+/-*^) but not wild-type mice in vivo. Two-photon calcium imaging in brain slice demonstrated profound impairment in filtering of perforant path input by DG in young adult *Scn1a*^*+/-*^ mice due to enhanced excitatory input to granule cells. Excitability of parvalbumin interneurons (PV-INs) was near-normal and selective activation of PV-INs rescued circuit impairments. This demonstrates developmental reorganization of hippocampal circuitry that can be modulated by recruitment of functional PV-INs, suggesting potential therapeutic approaches towards seizure modulation. The identified circuit abnormality mirrors that seen in models of chronic temporal lobe epilepsy, suggesting convergent mechanisms linking genetic and acquired causes of temporal lobe-onset seizures.

## Introduction

Pathogenic variants in *SCN1A*, which encodes the voltage-gated sodium channel *α* subunit Nav1.1, cause a spectrum of epilepsies including Dravet syndrome (DS)(***Claes et al., 2001***), a developmental and epileptic encephalopathy. DS patients exhibit refractory infantile-onset epilepsy with developmental delay and features of autism spectrum disorder, leading to severe and enduring cognitive impairment.

Most cases of DS are due to pathogenic loss of function variants in the *SCN1A* gene, leading to haploinsufficiency of Nav1.1. The heterozygous *Scn1a* mutant (*Scn1a*^*+/-*^) mouse is a well-established preclinical model of DS that recapitulates key phenotypic features of the human condition(***Mistry et al., 2014***). When expressed on a 50:50 129S6:C57BL6/J genetic background, these mice exhibit spontaneous seizures beginning at approximately post-natal day (P) 18, and high rates of sudden unexpected death in epilepsy (SUDEP) (***Mistry et al., 2014***). These mice also exhibit temperature-sensitive seizures, akin to seizures triggered in the setting of fever or hyperthermia in human patients with DS, which represents a key experimental advantage of this mouse model as it readily facilitates experiments on inducible but naturalistic seizures in vivo (***Tran et al., 2020***).

The prevailing theory as to how reduction in sodium current leads to epilepsy in DS is the so-called “interneuron hypothesis”, which posits that haploinsufficiency of Nav1.1 results in selective deficits in inhibitory interneuron excitability based on the relative reliance of this cell class on Nav1.1 for action potential generation. Electrophysiological recordings from acutely dissociated hippocampal neurons from *Scn1a*^*+/-*^ mice initially found decreased sodium current density selectively in bipolar-shaped presumptive GABAergic interneurons, but not in pyramidal cells (***Bechi et al., 2012; Yu et al., 2006***). Nav1.1 has been shown to be prominently expressed in parvalbumin-expressing GABAergic inhibitory interneurons (PV-INs) in neocortex (***Ogiwara et al., 2007***), as well as by somatostatin (SST-INs) and vasoactive intestinal peptide-expressing interneurons (VIP-INs). Consistent with these findings, impaired excitability of inhibitory interneurons – PV-INs as well as SST and VIP-INs – has been demonstrated in *Scn1a*^*+/-*^ mice (***Ogiwara et al., 2007; Tai et al., 2014; Favero et al., 2018; Goff and Goldberg, 2019***). This “interneuron hypothesis” is further supported by the fact that the DS phenotype is recapitulated by selective loss of Nav1.1 exclusively in GABAergic interneurons (***Cheah et al., 2012; Dutton et al., 2013; Rubinstein et al., 2015; Ogiwara et al., 2013***).

However, multiple lines of evidence suggest that the pathophysiology of epilepsy in *Scn1a*^*+/-*^ mice is more complex than impairment of inhibition. First, the VIP-INs are disinhibitory, so the hypoexcitability of this population (***Goff and Goldberg, 2019***) should in fact increase inhibition in neocortical circuits. Second, data from the *Scn1a*^*+/-*^ mouse models (***Mistry et al., 2014***) and from human-derived induced pluripotent stem cells (iPSCs) (***Jiao et al., 2013; Liu et al., 2013***) suggest that excitatory neurons in DS may also be hyperexcitable, although there remains controversy in the iPSC literature (***Sun et al., 2016***). Third, recent work found that impaired action potential generation present at early developmental time points in neocortical PV-INs normalized by P35 (***Favero et al., 2018***), whereas DS mice continue to exhibit epilepsy, cognitive impairment, and SUDEP. Finally, several in vivo studies have failed to find a decrease in interneuron activity in *Scn1a*^*+/-*^ mice (***Tran et al., 2020; De Stasi et al., 2016***).

Mechanisms of seizure generation and maintenance of chronic epilepsy remain unclear despite this now growing literature characterizing deficits in *Scn1a*^*+/-*^ mice at a single cell level (***Ogiwara et al., 2007; Tai et al., 2014; Favero et al., 2018; Goff and Goldberg, 2019; Cheah et al., 2012; Dutton et al., 2013***). This may be in part due to the lack of data linking cellular deficits to circuit-level abnormalities. Convergent data suggest that the dentate gyrus (DG) may be a key locus of pathology and seizure generation in *Scn1a*^*+/-*^ mice. Although *Scn1a*^*+/-*^ mice exhibit multifocal epilepsy, temperature-induced seizures have been shown to prominently emanate from the temporal lobe (***Liautard et al., 2013***), and focal hippocampal Nav1.1 reduction is sufficient to confer temperature-sensitive seizure susceptibility in conditional *Scn1a*^*+/-*^ mice (***Stein et al., 2019***). Although the DG receives a strong excitatory input from entorhinal cortex (***Amaral et al., 2007***), inhibitory micro-circuits within the DG regulate population activity such that granule cells (GCs) are only sparsely activated under physiologic conditions both in reduced preparations in vitro and in experimental animals in vivo (***Chawla et al., 2005; Diamantaki et al., 2016; Senzai and Buzsáki, 2017; Neunuebel and Knierim, 2012***). However, seizure-evoked immediate early gene activation is most apparent in the DG granule cell layer in *Scn1a*^*+/-*^ mice (***Dutton et al., 2017***), suggesting a circuit-level hyperexcitability in contrast to the normal state of the DG.

In this study, we demonstrate a profound hyperexcitability of the cortico-hippocampal circuit in young adult *Scn1a*^*+/-*^ mice that is not present at epilepsy onset using two-photon calcium imaging and cellular and synaptic physiology in an acute slice preparation. We find this circuit dysfunction is likely driven at least in part by excessive excitation, as opposed to being exclusively attributable to impaired inhibition. We then extend these findings *in vivo* using optogenetics in awake, behaving mice during temperature-sensitive seizures to demonstrate a role for this cortico-hippocampal dysregulation in ictogenesis. Finally, we recruit functional PV-INs in DG via selective optogenetic activation to rescue this circuit hyperexcitability in acute slice. These findings suggest that the cortico-hippocampal circuit may be a critical locus of pathology in DS and across divergent causes of temporal lobe-onset seizures.

## Results

### Response of dentate gyrus to entorhinal cortical input is selectively impaired in young adult *Scn1a*^*+/-*^ mice

Emerging data suggests an important role for the hippocampus and, more specifically, the dentate gyrus (DG), in Dravet syndrome (DS) pathology (***Dutton et al., 2013; Liautard et al., 2013; Stein et al., 2019***). We therefore hypothesized that the circuit function of DG – filtering input from the entorhinal cortex – might be impaired in *Scn1a*^*+/-*^ mice. We used two photon calcium imaging (2P imaging) to achieve large-scale quantification of granule cell (GC) activation in response to stimulation of the perforant path (PP), excitatory projections from entorhinal that are the major input to GCs. We first injected an adeno-associated virus (AAV) encoding the calcium indicator, GCaMP7s (***Dana et al., 2019***), under control of a pan-neuronal promoter, hSyn1, into DG of *Scn1a*^*+/-*^ mice and age-matched wild-type littermate controls (Figure 1A). We then prepared acute hippocampal-entorhinal cortex (HEC) slices (***Xiong et al., 2017***), which preserve the PP projection from entorhinal cortex to hippocampus within the slice (Figure 1B). We then performed 2P imaging of evoked calcium transients in cells within the granule cell layer (GCL) in response to electrical stimulation of the PP (Figure 1C-D).

**Figure 1.**
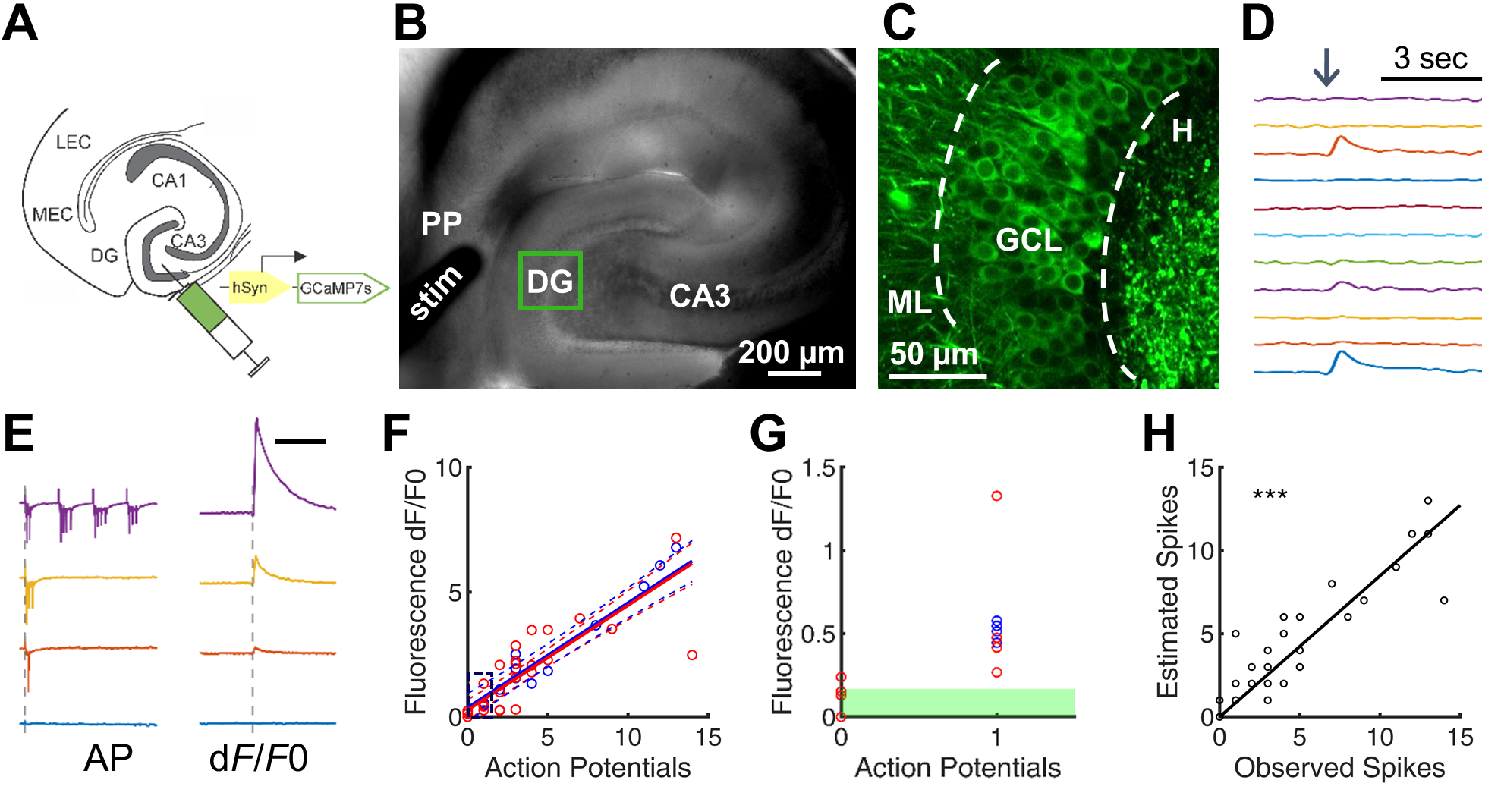
Two-photon calcium imaging of perforant path-evoked dentate gyrus activation in acute brain slice. (A) Mice were injected with AAV9-hSyn-GCaMP7s into DG. (B) Acute slices are cut at a 15° angle off-axial to maximize connectivity. Shown is the location of a stimulation electrode (stim) in the perforant path (PP) and the imaging field is in DG. (C) GCaMP labeling of GCs in the granule cell layer (GCL; dashed lines) between the hilus (H) and molecular layer (ML). (D) Example GC response to PP stimulation (arrow), displayed as change in fluorescence over baseline (dF/F0). (E) Representative data from a single GC firing 0, 1, 3, or 13 action potentials (left) in response to PP stimulation, while calcium transients (right) were simultaneously recorded. Scale bar represents 75 ms (left traces) and 5 s (right traces). Vertical dashed line indicates stimulus onset. (F) Action potentials versus calcium transient magnitude for GCs from WT (blue) and *Scn1a*^*+/-*^ (red) mice. Data were fit using a linear model, with no significant difference between genotypes (solid / dashed line = best-fit / 95% confidence interval). Young adult mice (P61-77); n = 7 cells from 5 mice. (G) Results for 0 and 1 action potentials (i.e. the region outlined in F). Threshold for action potential detection (green bar) was defined as p = 0.001 from a normal distribution fit to the dF/F0 values for 0 action potentials. (H) Observed action potentials versus action potentials derived from deconvolution (R^2^ = 0.83; p < 0.0001).

To determine the extent to which GCaMP7s reliably reports single action potentials in GCs under these experimental conditions, we performed simultaneous cell-attached recording and calcium imaging of GCaMP-expressing cells within the GCL in young adult (P61-81) *Scn1a*^*+/-*^ and wild-type mice. We used a quantum dot-labeled pipette tip (***Andrásfalvy et al., 2014***) for 2P-guided targeted recording. This allowed us to correlate calcium transients within individual cells against a gold-standard quantitative measure of action potentials (APs). We found that single action potentials were reliably detected Figure 1E-F), and that the number of action potentials and the magnitude of the calcium signal were highly correlated (R^2^ = 0.769). This linear relationship between AP and calcium signal was the same in wild-type and *Scn1a*^*+/-*^ mice across the measured range (0-14 APs; Figure 1F), indicating that GCaMP7s reports action potentials in GCs similarly in both genotypes and validating 2P calcium imaging as a method for comparing large-scale DG excitability between wild-type and *Scn1a*^*+/-*^. To confirm that GCaMP7s was adequately sensitive to detect single action potentials across tens to hundreds of cells, we fit a normal distribution to the dF/F0 values associated with 0 action potentials, set a threshold defined by this data (p = 0.001), and verified that 100% of single APs recorded resulted in signal above that threshold (n = 8 action potentials from 6 cells; Figure 1G). Finally, we deconvolved the dF/F0 signals (***Pnevmatikakis et al., 2016***) to extract the number of action potentials from the calcium transients and verified the deconvolution algorithm on the “ground truth” dataset (R^2^ = 0.83; p < 0.0001; Figure 1H).

In order to quantify the extent of DG activation in response to entorhinal cortex input, we stimulated the PP while performing 2P imaging of the evoked responses in GCs (Figure 2). We tested early postnatal (P14-21; at/around epilepsy onset) and young adult (P48-91; chronic phase) mice. We delivered either a single pulse or a train of 4 pulses at 20 Hz across a range of stimulation intensities, within the standard range of paradigms used in the field (***Yu et al., 2013; Ewell and Jones, 2010; Dengler et al., 2017***). We then identified activated GCs within the imaging field in response to each stimulation condition, defining activation as an increase in calcium signal (dF/F0) at least 3 times the standard deviation above the mean noise (Figure 2, columns 1-4) or if one or more action potentials were extracted using deconvolution (Figure 2, column 5). To determine if the responses across stimulation intensities differed by genotype, we employed a mixed model statistical approach, to account for potential variation between individual imaging fields, slices, and/or mice.

**Figure 2.**
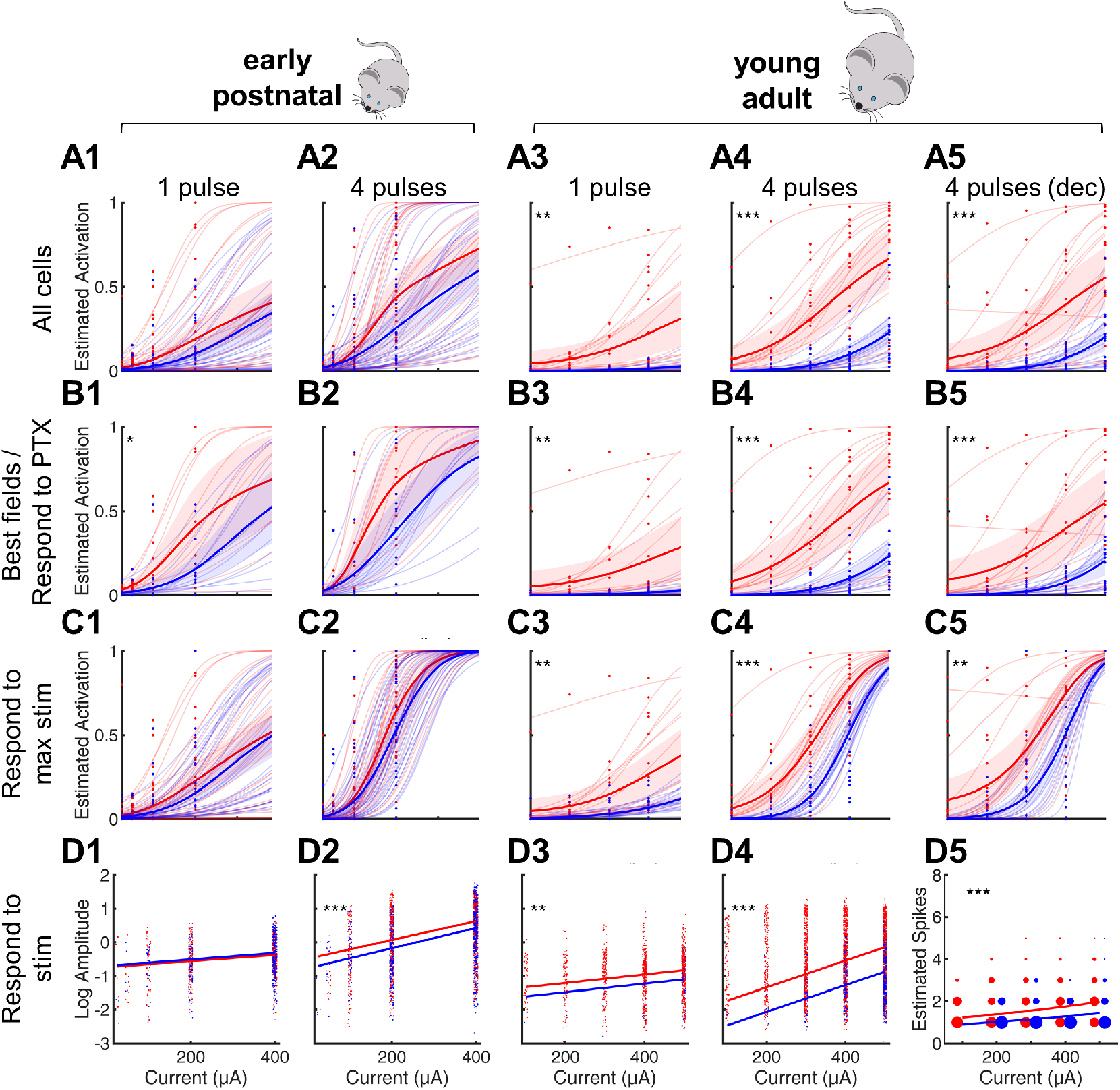
Selective impairment of dentate gyrus function in young adult *Scn1a*^*+/-*^ mice. Proportion of activated GCs. (A-C) and magnitude of activation (D) in response to PP stimulation (early postnatal (P14-21; columns 1 and 2) and young adult (P48-91; columns 3 and 4), with deconvolved data for the young adult time point shown in column 5. Wild-type in blue; *Scn1a*^*+/-*^ in red. Data were analyzed using a mixed model to account for potential variability between animal, slice, field, and cell. Proportion of responsive GCs was calculated relative to all GCaMP-expressing cells (A), restricting analysis to either the most responsive 2 fields per mouse (for early postnatal mice; B1, B2) or to only GCs that respond to PP stimulation in the presence of 100 µM picrotoxin (for young adult mice; B3-5), or restricting analysis to only GCs that respond to the maximal stimulation delivered (C). (D) Magnitude of activated GC responses to PP stimulation, expressed as natural log dF/F0 (columns 1-4) or estimated spikes based on deconvolution (column 5; size of data points reflects number of cells at that value). Dots: raw data from all imaging fields or cells; light lines: predicted fits for each field; dark lines: average of fits; shaded: 95% confidence intervals of average fits. Stars indicate significant differences in curve fits: * denotes p < 0.05, ** denotes p < 0.01, and *** denotes p < 0.001. n for early postnatal mice (P14-21; *Scn1a*^*+/-*^, WT) = 5,6 (mice); 12, 15 (slices); 31,35 (fields); 765, 619 (cells). n for young adult mice (P48-91; *Scn1a*^*+/-*^, WT) = 8,6 (mice); 17,17 (slices and fields); 1236, 1167 (cells).

We first calculated the proportion of activated GCs relative to all identifiable GCaMP7s-expressing cells, fit the data from each genotype (binomial distribution fitted with a probit link function), and compared the fit of the curves between genotypes (“All cells” in Figure 2A). We found no statistically significant difference between genotypes at the early postnatal timepoint (Figure 2A, columns 1-2). At the young adult timepoint, however, a significantly higher proportion of GCs were activated by PP input in *Scn1a*^*+/-*^ mice than in age-matched wild-type controls (p < 0.001; Figure 2A, columns 3-5). For instance, *Scn1a*^*+/-*^ GC activation was over 3-fold larger than that of wild-type (0.508 ± 0.078 versus 0.161 ± 0.002; p < 0.001) in response to 4 × 400 µA pulses (subset of data in graph A1).

We validated these data in several different ways to account for the possibility that some GCs might be unresponsive due to deafferentation during the slice preparation. First, for early postnatal mice, as data was obtained from multiple imaging fields per slice, we reasoned that the fields with the highest proportion of responsive cells were likely the best anatomically connected, so we performed a subgroup analysis restricted to the two most responsive imaging fields per mouse (“Best fields” in Figure 2B, columns 1-2). We found that response of DG in early postnatal *Scn1a*^*+/-*^ mice to the single pulse stimulation condition did reach statistical significance in this case (p < 0.05) although the response to 4 pulses did not. Second, for the young adult mice, in which we imaged only one field per slice, we re-imaged each field in the presence of picrotoxin (PTX), a GABA_A_ receptor antagonist that is known to induce widespread PP-driven recruitment of GCs (***Dengler et al., 2017***). We then excluded from analysis any GC that failed to respond to PP stimulation in the presence of PTX (“Respond to PTX” in Figure 2B, columns 3-5), and re-demonstrated highly significant *Scn1a*^*+/-*^ GC hyperactivation in young adult mice. Third, for all data, we repeated the analysis to include only the GCs that responded to the highest amplitude (400 or 500 µA) PP stimulation delivered (“Responsive to max stim” in Figure 2C), and found highly significant *Scn1a*^*+/-*^ GC hyperactivation for the young adult, but not early postnatal, datasets. Results based either on the calculated dF/F0 value, and those obtained via deconvolution, were similar (Figure 2A-C, column 4 vs. 5).

Finally, we quantified the magnitude of the evoked calcium signal (dF/F0) and used a mixed model approach to compare across genotypes, using a normal distribution fitted with a logarithmic link function (Figure 2D, Columns 1-4). We included only activated GCs in this analysis (“Responds to max stim”). For early postnatal mice, the response magnitude of activated GCs was the same between wild-type and *Scn1a*^*+/-*^ mice in the single pulse condition; for the 4 pulse condition, there was a small (22%) increase in response magnitude in GCs from *Scn1a*^*+/-*^ mice. In young adult mice, the *Scn1a*^*+/-*^ GC response magnitude was larger than wild-type in both stimulation conditions, and the effect size was much larger (130% increase) in the 4 pulse condition. The same result was obtained using estimated action potential numbers calculated via deconvolution (Figure 2D, column 5).

Overall, these findings demonstrate a profound impairment of the cortico-hippocampal circuit selectively in young adult *Scn1a*^*+/-*^ mice, with PP stimulation resulting in greater proportional activation of GCs, as well as an increase in the number of action potentials fired in those GCs that were activated. Results were highly significant and robust to various analyses at the young adult time point, suggesting that circuit dysfunction develops over or is unmasked with time.

### Mechanisms of dentate gyrus circuit dysfunction in young adult *Scn1a*^*+/-*^ mice

Increased PP activation of GCs could be due to increased intrinsic GC excitability, dysfunction of feed-forward inhibition, or increased synaptic excitatory drive onto GCs. To investigate the mechanism of DG circuit dysfunction in young adult *Scn1a*^*+/-*^ mice, we first assessed the intrinsic excitability of GCs in *Scn1a*^*+/-*^ mice versus controls. Whole-cell current-clamp recordings of GCs from *Scn1a*^*+/-*^ mice and age-matched wild-type littermate controls (P35-77) showed no statistical differences across a range of measures of intrinsic excitability, properties of individual action potentials (APs), and repetitive AP firing (Table 1).

**Table 1.** Properties of dentate gyrus granule cells from *Scn1a*^*+/-*^ and wild-type mice. All statistical comparisons are made considering the average of each mouse as n = 1. Note that the p values listed have not been corrected for multiple comparisons, but remain non-significant. Young adult mice (P35-77); n = 7 mice (16 cells) for *Scn1a*^*+/-*^ and 5 mice (24 cells) for wild-type.

| Measurement | *Scn1a*<sup>+/-</sup><br>n = 7 mice (16 cells) | WT<br>n = 5 mice (24 cells) | p-value |
| --- | --- | --- | --- |
| V<sub>m</sub> (mV) | -79.8 ± 1.1 | -78.2 ± 1.8 | 0.45 |
| R<sub>m</sub> (MΩ) | 353 ± 35 | 396 ± 64 | 0.54 |
| Rheobase (pA) | 87 ± 11 | 86 ± 19 | 0.98 |
| 1<sup>st</sup> AP peak (mV) | 45.3 ± 1.5 | 42.8 ± 3.6 | 0.48 |
| AP half width (ms) | 0.90 ± 0.02 | 0.88 ± 0.08 | 0.84 |
| Max instantaneous (Hz) | 191 ± 12 | 179 ± 8 | 0.47 |
| Max steady-State (Hz) | 64 ± 5 | 53 ± 2 | 0.13 |

Single-cell electrophysiology data from acute brain slices and acutely dissociated neurons prepared from various *Scn1a*^*+/-*^ mouse lines from multiple laboratories has repeatedly identified interneuron dysfunction (***Ogiwara et al., 2007; Tai et al., 2014; Cheah et al., 2012; Rubinstein et al., 2015; Dutton et al., 2017; Richards et al., 2018***) and in particular dysfunction of parvalbumin-expressing fast-spiking GABAergic interneurons (PV-INs) (***Tai et al., 2014; Dutton et al., 2013; Ru-binstein et al., 2015***). Therefore, we next tested the intrinsic properties of DG PV-INs in *Scn1a*^*+/-*^ mice versus wild-type littermate controls based on the involvement of these cells in DS pathogenesis as well as the fact that the GC response to PP input is known to be powerfully regulated by feedforward inhibition mediated by DG PV-INs (***Lee et al., 2016***).

We prepared acute brain slices from *Scn1a*^*+/-*^ mice and wild-type littermates (P61-77) expressing tdTomato (tdT) under PV-specific Cre-dependent control, as described previously (***Favero et al., 2018***). PV-INs in DG were thus identified by endogenous tdT expression visualized with epifluorescence and characteristic location at the GCL:hilus border. Data obtained from whole-cell patch clamp recordings of these cells showed no statistical difference across measures of intrinsic excitability, properties of individual action potentials, and maximal firing frequencies (Table 2). *Scn1a*^*+/-*^ PV-INs fired normally at the onset of a depolarizing current step and reached identical maximal instantaneous and steady-state firing frequencies. We did identify diverging responses during sustained trains, with *Scn1a*^*+/-*^ PV-INs exhibiting gradual spike height accommodation (Figure 3A-C); this ultimately progresses to spike failure with prolonged and large-amplitude (2.5-fold rheobase) current injections (Figure 3D-E).

**Table 2.**
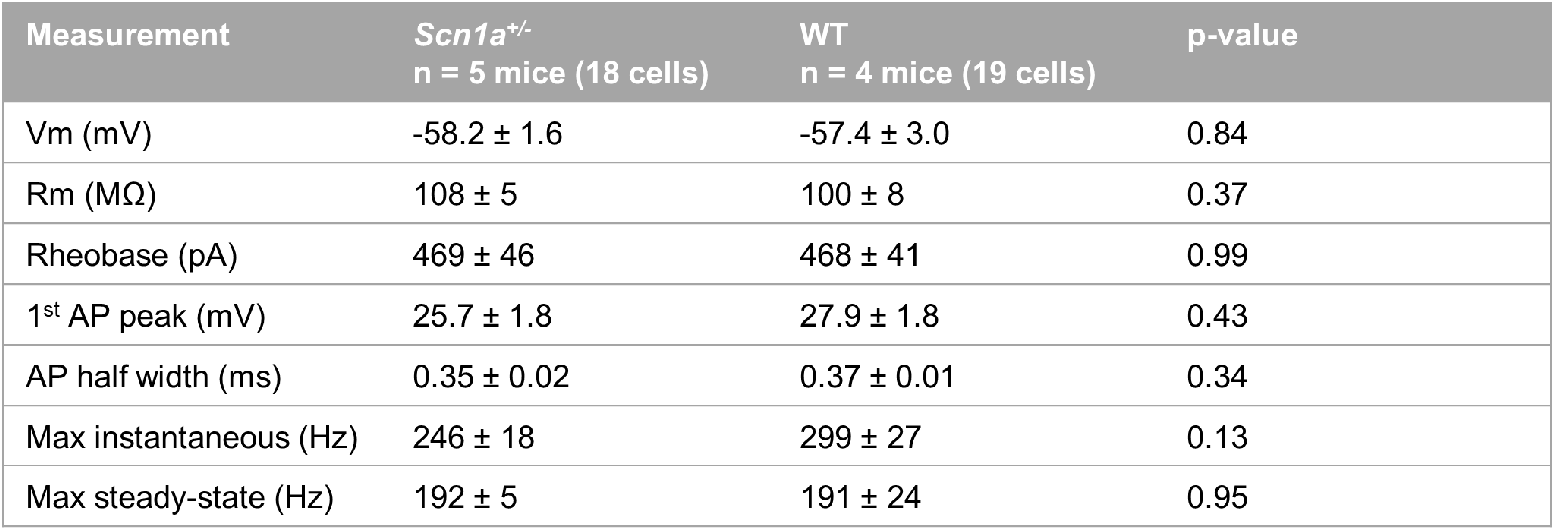
Properties of DG PV-INs from *Scn1a*^*+/-*^ and wild-type mice. All statistical comparisons are made considering the average of each mouse as n = 1. Note that the p values listed have not been corrected for multiple comparisons, but remain non-significant. Young adult mice (P61-77); n = 5 mice (18 cells) for *Scn1a*^*+/-*^ and 4 mice (19 cells) for wild-type.

| Measurement | *Scn1a*<sup>+/-</sup><br>n = 5 mice (18 cells) | WT<br>n = 4 mice (19 cells) | p-value |
| --- | --- | --- | --- |
| V<sub>m</sub> (mV) | -58.2 ± 1.6 | -57.4 ± 3.0 | 0.84 |
| R<sub>m</sub> (MΩ) | 108 ± 5 | 100 ± 8 | 0.37 |
| Rheobase (pA) | 469 ± 46 | 468 ± 41 | 0.99 |
| 1<sup>st</sup> AP peak (mV) | 25.7 ± 1.8 | 27.9 ± 1.8 | 0.43 |
| AP half width (ms) | 0.35 ± 0.02 | 0.37 ± 0.01 | 0.34 |
| Max instantaneous (Hz) | 246 ± 18 | 299 ± 27 | 0.13 |
| Max steady-state (Hz) | 192 ± 5 | 191 ± 24 | 0.95 |

**Figure 3.**
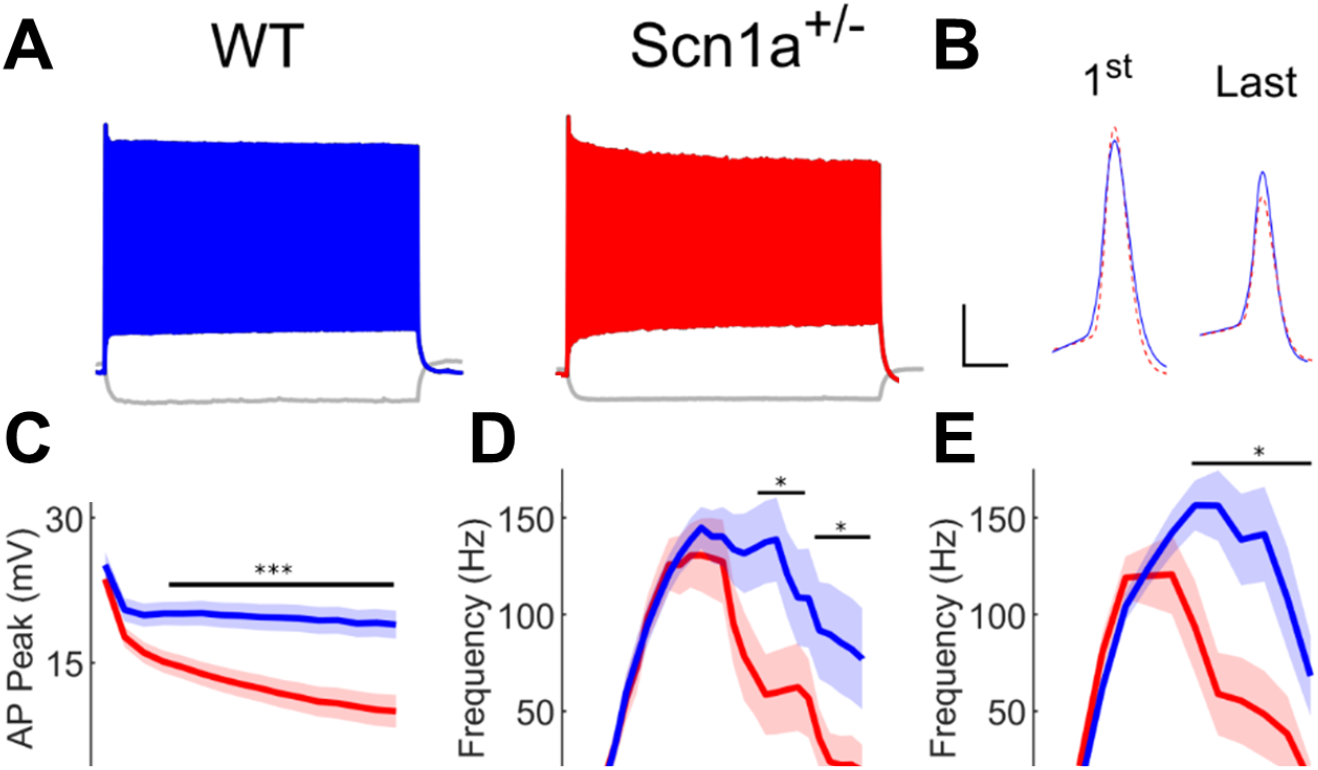
Subtle impairment of spike generation in *Scn1a*^*+/-*^ DG PV-INs. (A) Current clamp recordings from WT (blue) and *Scn1a*^*+/-*^ (red) DG PV-INs, illustrating a progressive spike-height accommodation seen in *Scn1a*^*+/-*^ PV-INs in response to a prolonged depolarizing current step (show at 3X rheobase). Scale bar 20 mV / 100 ms. (B) Example individual superimposed WT and *Scn1a*^*+/-*^ action potentials (APs) from the beginning (1st) and end (last) of a 600 ms depolarizing pulse train. Note that the 1st AP morphology is near-identical whereas the last *Scn1a*^*+/-*^ AP has a lower peak amplitude and spike height. Scale bar 20 mV / 1 ms. (C) AP peak as a function of AP number. (D) Current/frequency (I-F) plot for GC PV-INs. (E) I-F plot with current normalized to Rheobase for each cell. Line and shaded areas represent mean and SEM. Bars indicate significance calculated using one-way ANOVA and post-hoc tests with Bonferroni correction; * represents < 0.05 and *** represents < 0.001. Young adult mice (P61-77); n = 4 mice (19 cells) for wild-type and 5 mice (18 cells) for *Scn1a*^*+/-*^.

To further confirm that the deficits seen in individual PV-IN firing properties (Figure 2) were not responsible for the GC hyperactivation observed in *Scn1a*^*+/-*^ mice, we performed an additional set of experiments using Hm1a, a peptide toxin that acts as an Nav1.1-specific activator (***Osteen et al., 2016***) and has been shown to correct the abnormalities seen in PV-INs in *Scn1a*^*+/-*^ mice (***Goff and Goldberg, 2019***). We found that bath-application of Hm1a decreased PV-IN spike failure in response to large and prolonged current injections (Figure 4A-B). However, this did not result in a decrease in large-scale GC activation as measured via 2P imaging; i.e. there was no significant change between responses at baseline versus after Hm1a wash-in for any of the multiple stimulation conditions tested. PP stimulation of 4 pulses at 500 µA (Figure 4C) produced no change in the net dF/F0 measured globally across the GCL (dF/F0 = 0.63 ± 0.17 at baseline and 0.58 ± 0.17 post Hm1a; p = 0.52; paired t-test). This result suggests that the subtle identified deficits in PV-IN spike generation and impairment in repetitive firing does not underlie the observed large-scale circuit impairment.

**Figure 4.**
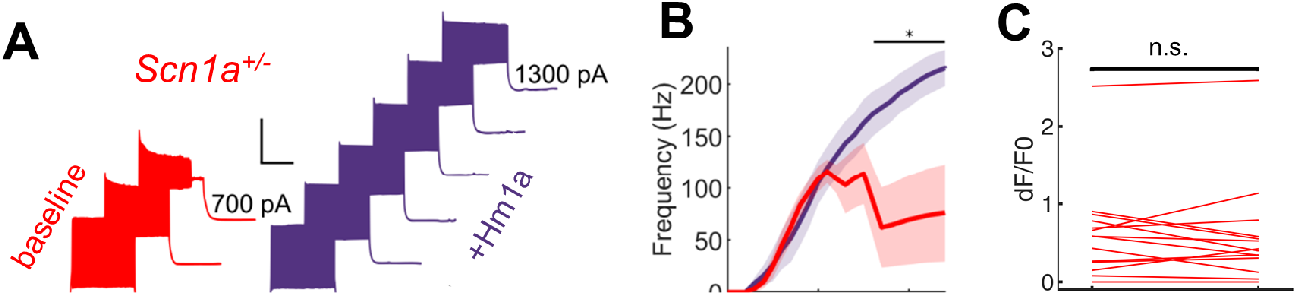
Hm1a enhances *Scn1a*^*+/-*^ DG PV-IN fast-spiking firing properties but does not rescue evoked granule cell activation. Example traces from one *Scn1a*^*+/-*^ PV-IN (A) as well as population data (B) showing within-cell comparisons of responses to depolarizing current steps at baseline (red) and in the presence of Hm1a (purple). Hm1a decreased spike height accommodation and prevented AP failures with larger current injections. Line and shaded areas represent mean and SEM. Bars indicate significance calculated using one-way ANOVA and post-hoc tests with Bonferroni correction; * represents < 0.05. (C) 2P calcium imaging data with bulk GC response (dF/F0) to 4 pulses at 400 µA PP stimulation, using the entire granule cell layer as a Region of Interest, recorded within the same imaging fields at baseline and after a 20-25 minute wash-on of Hm1a. Hm1a did not change the GC response to PP stimulation (dF/F0 = 0.63 ± 0.17 at baseline and 0.58 ± 0.17 post Hm1a; p = 0.52; paired t-test). n = 3 mice (5 cells) for PV-IN patching data and 7 mice (14 imaging fields) for 2P imaging.

Since impaired response of DG to entorhinal cortex input in *Scn1a*^*+/-*^ mice could not be adequately attributed to aberrant intrinsic properties of either DG GCs or PV-INs, we next considered whether this finding may instead be due to alterations in the excitatory drive onto GCs and/or recruitment of disynaptic inhibition. We performed whole-cell voltage-clamp recordings in GCs from *Scn1a*^*+/-*^ mice versus wild-type controls (P54-75) using a cesium-based internal solution to isolate evoked monosynaptic excitatory postsynaptic currents (EPSCs; recorded at -70 mV) and di-synaptic inhibitory postsynaptic currents (IPSCs; recorded at +10 mV) (Figure 5A). We first quantified the minimal PP stimulation required to evoke an EPSC and IPSC for each cell (Figure 5B) and found no significant difference between genotypes.

**Figure 5.**
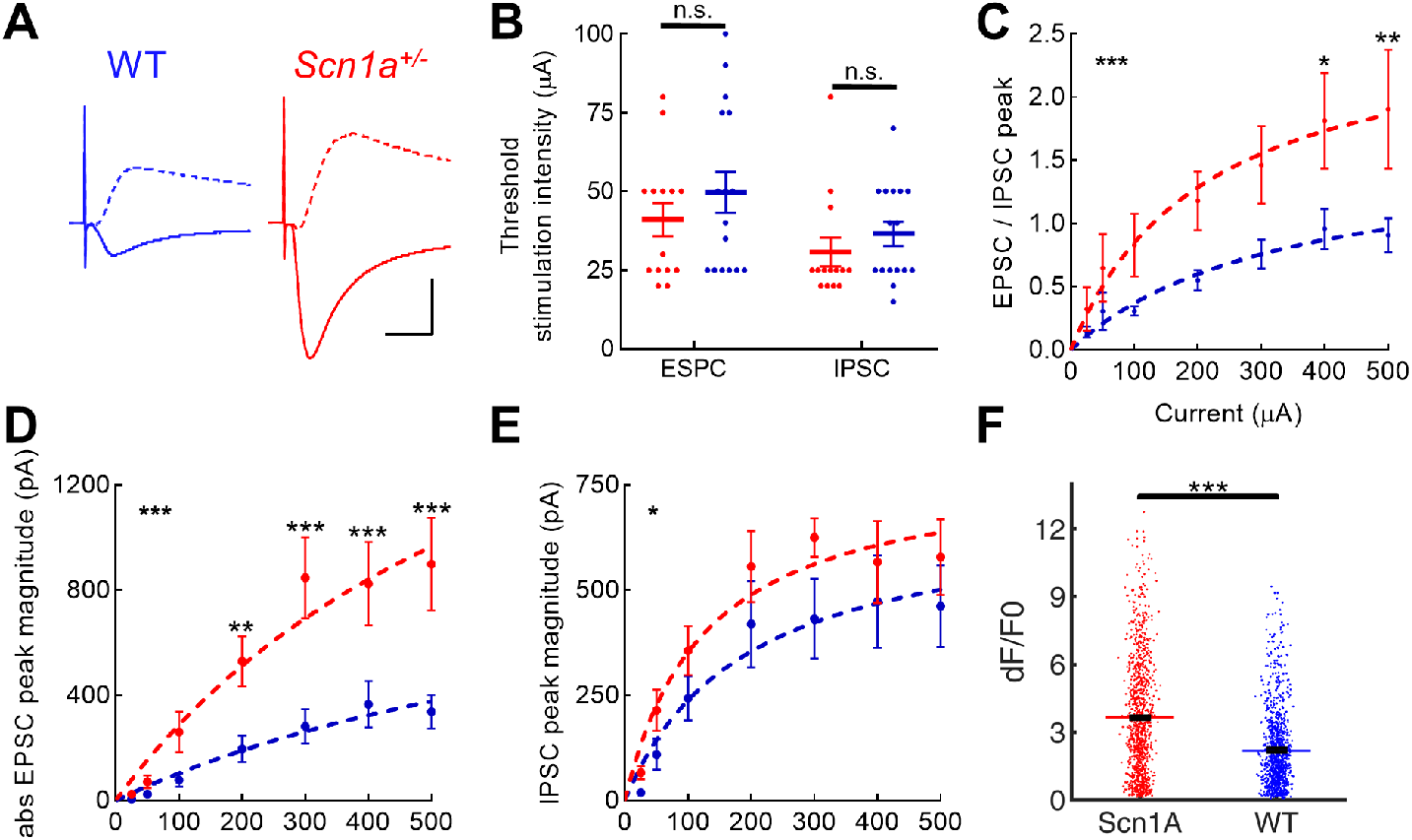
Selective increase in PP-evoked excitation of *Scn1a*^*+/-*^ dentate gyrus granule cells. Whole cell patch clamp recordings were performed in GCs from *Scn1a*^*+/-*^ and wild-type mice (P54-75). (A) Representative traces from wild-type (blue) and *Scn1a*^*+/-*^ (red) GCs showing evoked monosynaptic EPSC (recorded at -70 mV; solid) and di-synaptic IPSCs (recorded at +10 mV; dashed) in response to 300 µA PP stimulation. Scale bar, 10 ms / 500 pA. (B) The threshold EPSC and IPSC stimulation intensity was identified for each cell, with no significantly difference between genotypes. (C) EPSC / IPSC magnitude was calculated in response to 100-500 µA PP input. Data were fit with a one site binding curve and compared with an extra sum-of-squares F test (p < 0.0001). Mixed-effects analysis with multiple comparisons were used for comparisons at each point (p = 0.025 for 400 µA and p = 0.008 for 500 µA). (D) Raw evoked ESPC magnitude was significantly larger in *Scn1a*^*+/-*^ GCs (p < 0.0001; extra sum-of-squares F test comparison of curve fits). Mixed-effects analysis with multiple comparisons were used for comparisons at each point (p = 0.005 for 200 µA and p < 0.0001 for 300-500 µA). (E) Raw evoked disynaptic IPSC magnitude was also higher overall in *Scn1a*^*+/-*^ GCs (p = 0.02, extra sum-of-squares F test comparison of curve fits), although significance was not reached at any individual data point. (F) Magnitude of evoked GC responses as quantified by calcium imaging (dF/F0) in the presence of 100 µM picrotoxin, with significantly larger responses measured in *Scn1a*^*+/-*^ GCs despite GABA_A_ receptor blockade (p <0.001; mixed model analysis). For patching data, n = 14 cells (4 mice) for *Scn1a*^*+/-*^ and n = 16 cells (4 mice) for wild-type. For 2P imaging data, n (*Scn1a*^*+/-*^, WT) = 7,6 (mice); 14,16 (slices and fields); 1073, 1109 (cells). For C-E, stars in the upper left indicate significance of overall curve fits while stars above individual data points indicate post-hoc significance at each point.

We next calculated the ratio of the maximal evoked EPSC/IPSC amplitude (Figure 5C). We tested the null hypothesis that all data could be fit by one curve (one-site binding curve), and found that, instead, the fits of the two genotypes were highly statistically different (p < 0.0001), with a markedly larger EPSC/IPSC ratio in *Scn1a*^*+/-*^ GCs. The data were further analyzed using a mixed-effects posthoc analysis with multiple comparisons, with significant differences found between genotypes in the two strongest stimulation conditions (400 µA: 1.8 ± 0.4 versus 1.0 ± 0.2, p = 0.045; 500 µA: 1.9 ± 0.5 versus 1.0 ± 0.1, p = 0.008).

To understand whether this result was driven by increased excitation and/or decreased inhibition, we examined the raw EPSC and IPSC magnitudes. We found significantly larger EPSC magnitudes (Figure 5D) in *Scn1a*^*+/-*^ versus wild-type GCs (p < 0.0001, analysis of curve fits), with over two-fold higher magnitudes in *Scn1a*^*+/-*^ (e.g., 400 µA: 825 ± 159 versus 365 ± 88 pA, p < 0.0001, using mixed-effects analysis with correction for multiple comparisons). Overall the IPSC magnitude also differed between genotypes (p = 0.02, analysis of curve fits), with IPSC amplitude trending larger (not smaller as would contribute to a larger E/I ratio) in *Scn1a*^*+/-*^ versus wild-type GCs, although the difference was not enough to reach significance at any individual data point on post-hoc analysis (Figure 5E).

Overall these data suggest that hyper-activation of GCs in young adult *Scn1a*^*+/-*^ mice is driven by increased excitatory drive to DG rather than impaired inhibition. If anything, disynaptic inhibition – mediated largely by PP-driven recruitment of PV-INs – is normal or enhanced, although unable to keep up with the greatly enhanced excitatory drive. To corroborate this finding, we further analyzed the calcium imaging data obtained in the presence of a saturating concentration of PTX. We postulated that if decreased GABA_A_ receptor-mediated inhibition was the cause of GC hyperactivation in *Scn1a*^*+/-*^ mice, then PTX should eliminate the difference between genotypes seen in Figure 2. However, we found that the genotype difference persisted in the presence of PTX, with mixed model analysis of dF/F0 magnitudes being larger for *Scn1a*^*+/-*^ (3.64 ± 0.07) versus wild-type (2.21 ± 0.10; p < 0.001) in the presence of PTX, further consistent with increased excitatory drive in *Scn1a*^*+/-*^ mice (Figure 5F).

### Seizures are triggered by optogenetic activation of entorhinal cortex in *Scn1a*^*+/-*^mice but not wild-type controls

The DG controls hippocampal signal propagation from entorhinal cortex to CA3; dysfunction of this control point is hypothesized to be involved in temporal lobe epileptogenesis and seizure generation in chronic temporal lobe epilepsy (***Dengler et al., 2017; Behr et al., 1998; Lu et al., 2016***). Having demonstrated that cortico-hippocampal circuit function is impaired in *Scn1a*^*+/-*^ mice (Figure 2), we hypothesized that stimulation of entorhinal cortex would engage hyperexcitable hippocampal circuits sufficient to induce seizures in vivo.

In preparation for moving in vivo, we first confirmed in slice that GCs could be efficiently activated not only by electrical stimulation (as above), but also via photostimulation of an excitatory opsin introduced into entorhinal cortex (Figure 6A-C). We then performed an in vivo experiment to test the implications of this circuit dysregulation on ictogenesis. We injected an AAV encoding ChR2 under control of the excitatory neuron-specific promoter, CaMKII, into the entorhinal cortex of *Scn1a*^*+/-*^ and wild-type mice. We then optogenetically stimulated entorhinal cortex in vivo, using parameters (5 ms pulse width at 20 Hz = 10% duty cycle; 5 seconds on / 5 seconds off) similar to previously published protocol (***Leung et al., 2018***). We observed that stimulation of entorhinal cor-tex was sufficient to trigger behavioral seizures (modified Racine scale (***Erum et al., 2019***) score 5) at room temperature in 4 out of a total of 9 *Scn1a*^*+/-*^ mice, whereas 0 out of 5 wild-type mice had seizures evoked by identical stimulation parameters.

**Figure 6.**
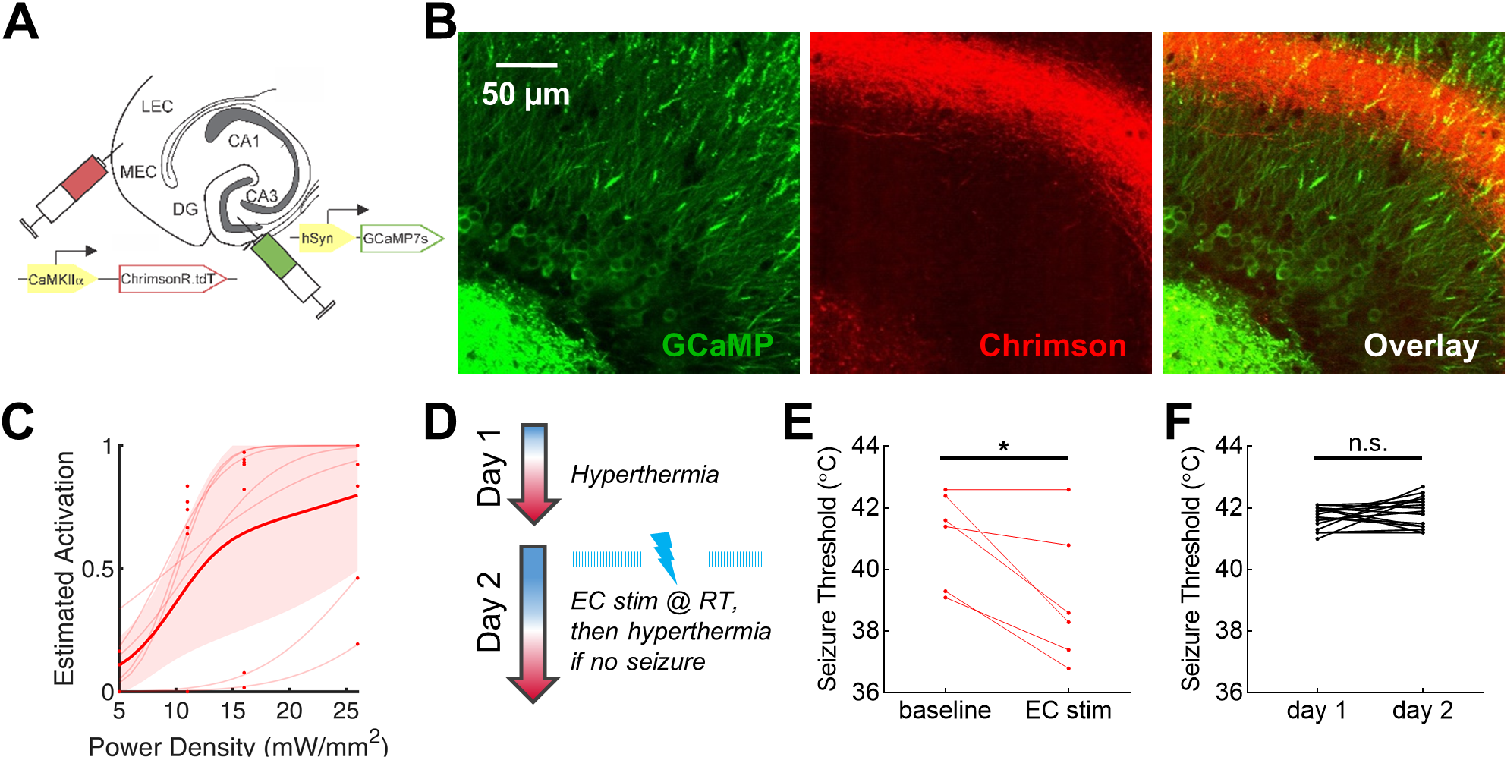
Activation of entorhinal cortex is ictogenic in *Scn1a*^*+/-*^ mice. (A) *Scn1a*^*+/-*^ mice were injected with both AAV9-hSyn-GCaMP7s into DG and AAV9-hSyn-ChrimsonR-tdT into EC. (B) Expression in the DG of GCaMP7s in granule cell bodies (green), and ChrimsonR-expressing fibers (red) projecting from entorhinal cortex to the DG molecular layer. (C) Robust response of GCs to photostimulation of fibers within the slice, across a range of power. n = 3 mice, 6 slices and fields, 1568 cells. (D) *Scn1a*^*+/-*^ mice were injected with ChR2 in entorhinal cortex. On Day 1, mice were heated to an internal body temperature of up to 42.5°C, or until a behavioral seizure was observed. On Day 2, pulsed photostimulation was delivered to EC at 20 Hz, 5 ms pulse-width, for 5 seconds on / 5 seconds off. Mice are stimulated initially at room temperature for up to 10 minutes, then while subjected to hyperthermia (again until 42.5°C, or until a behavioral seizure is observed). (E) Seizure threshold for *Scn1a*^*+/-*^ mice at baseline (Day 1) was significantly higher than when subjected to EC stimulation (Day 2) (41.1 ± 0.6 versus 39.1 ± 0.9; p = 0.02 via paired t-test; n = 6 mice). (F) Adult *Scn1a*^*+/-*^ mice were subject to hyperthermia on sequential days (separated by approximately 24 hours) until behavioral seizures were observed. There was no significant difference in seizure threshold across days (41.7 ± 0.1 versus 41.9 ± 0.1; p = 0.22 via paired t-test; n = 16 mice).

It is well established that *Scn1a*^*+/-*^ mice exhibit not only spontaneous but also temperature-sensitive seizures (***Oakley et al., 2009***). We took advantage of this phenotype to test whether activation of entorhinal cortex could also shift the threshold for seizure generation in *Scn1a*^*+/-*^ mice by performing a within-mouse comparison of seizure threshold, with or without EC stimulation. On Day 1, we elevated the body temperature of the mice in the absence of stimulation to determine baseline seizure threshold: all *Scn1a*^*+/-*^ mice (n = 6) had behavioral seizures at or prior to the end-point of 42.5°C body temperature, whereas no wild-type mice (n = 5) had behavioral seizures. On Day 2 (approximately 24 hours later), we optogenetically stimulated the mice, initially at room temperature (up to 10 minutes, or until a seizure was observed), and then (if no seizure), while subjecting the mice to hyperthermia (Figure 6D). The *Scn1a*^*+/-*^ mice exhibited behavioral seizures at a significantly lower threshold temperature (39.1 ± 0.9 versus 41.1 ± 0.6 °C; p = 0.02) with stimulation compared to baseline (Figure 6E). Only one wild-type mouse was observed to have a behavioral seizure during simultaneous optogenetic stimulation and heating (at 39.9°C internal body temperature), as has been previously observed in C57BL/6J mice (***van Gassen et al., 2008***).

We opted for this within-mouse experimental design to improve statistical power, but there is a theoretical concern that seizure threshold on Day 2 (the second consecutive day of stimulation) could be lowered by a seizure 24 hours prior (a “kindling”-like phenomenon). It has in fact been shown that, in young *Scn1a*^*+/-*^ mice (P18-19), “priming” mice (via a temperature-sensitive seizure) exacerbates the subsequent epilepsy phenotype (***Hawkins et al., 2017***). We tested seizure threshold on consecutive days in a separate cohort of mice (Figure 6F), and found that it was not significantly different (p = 0.22; n = 16 mice), suggesting that there is no equivalent priming effect in older mice (***Yamagata et al., 2020***).

### Rescue of dentate gyrus hyperexcitability by optogenetic activation of local PV-INs

Given that PV-INs are known to exert powerful feedforward inhibition in DG (***Lee et al., 2016***), yet we found DG PV-INs to be near-normal in *Scn1a*^*+/-*^ mice at the developmental time point examined (young adult; P63-74), we hypothesized that targeted activation of DG PV-INs could recruit additional GABAergic inhibition and correct the imbalance of inputs onto GCs and thereby rescue the over-activation of GCs by PP input in *Scn1a*^*+/-*^ mice. We injected a 1:1 mix of two AAVs into the DG of *Scn1a*^*+/-*^.PV-Cre mice (Figure 7A): (1) the calcium indicator GCaMP7s (AAV9.hSyn.GCaMP7s) as in Figure 1, and (2) the red-shifted excitatory opsin, ChrimsonR (***Klapoetke et al., 2014***) (AAV9.hSyn.FLEX.- ChrimsonR.tdT), allowing simultaneous imaging of DG and photo-activation of PV-INs (Figure 7B). Whereas GCaMP7s was expressed strongly in GCs, ChrimsonR expression was restricted to PV-INs, and ChrimsonR-expressing fibers were seen densely within the granule cell layer (Figure 7C), consistent with known perisomatic targeting of GCs by PV-INs (***Amaral et al., 2007; Houser, 2007***).

**Figure 7.**
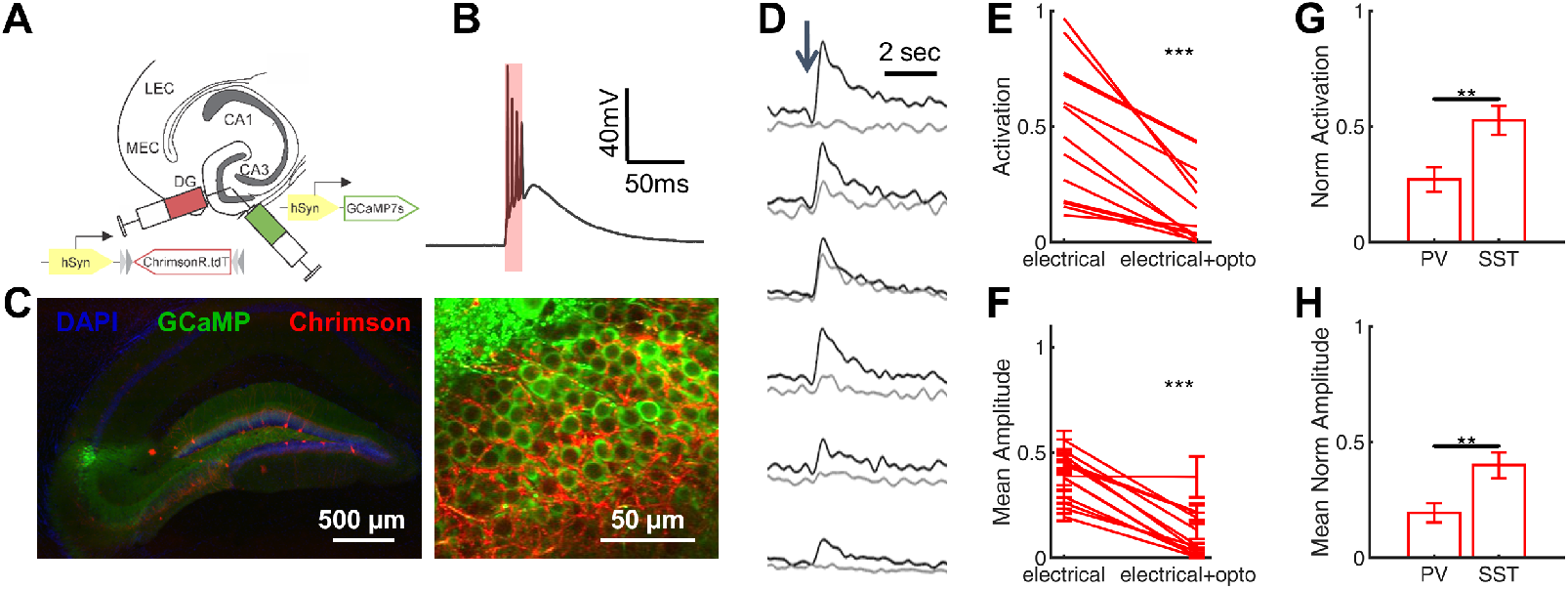
Rescue of cortico-hippocampal hyperexcitability via optogenetic activation of PV interneurons. (A) *Scn1a*^*+/-*^.PV-Cre mice were injected with a mix of AAV9-hSyn-GCaMP7s and AAV9-hSyn-Flex-ChrimsonR-tdT into DG. (B) Whole-cell current-clamp trace demonstrating light-evoked action potentials in a ChrimsonR-expressing PV cell (660nm, red bar). (C) Low and high magnification images showing the dentate gyrus (anatomically delineated by DAPI staining on the left, blue) with sparse PV cell bodies but dense PV fibers (red) surrounding GCaMP-expressing GCs (green) (D) Example calcium transients in GCs with electrical PP stimulation (single pulse, 500 µA, arrow), either alone (black) or coupled with optogenetic PV cell activation (10 ms light pulse; gray). Proportional GC activation (E) and magnitude of GC responses (F) to PP stimulation (1 pulse, 500 µA) are both significantly decreased by concurrent optogenetic PV cell activation. Each line represents one imaging field, subject to PP stimulation alone versus PP stimulation with optogenetic PV activation (p < 0.001; paired t-test). Optogenetic activation of DG PV-INs is more effective that activation of DG SST cells at decreasing PP stimulation-evoked activity in GCs, with a lower proportional activation (G) and dF/F0 amplitude (H), normalized to the response to PP stimulation alone in the same cells and imaging fields. Significance (p < 0.01) determined using mixed model analysis. PV-Cre mice: P61-71; n = 6 mice, 13 imaging fields, 1055 cells. SST-Cre mice: P68-71; n = 3 mice, 15 imaging fields, 663 cells.

We used 2P calcium imaging to record PP stimulation-evoked activation of GCs either with or without concurrent optogenetic activation of DG PV-INs. We found that optogenetic co-activation significantly reduced PP-evoked DG hyper-activation in *Scn1a*^*+/-*^ mice (Figure 7D-F), decreasing the proportion of activated GCs (0.15 ± 0.04 with PV-IN activation versus 0.48 ± 0.08 without; p < 0.001, paired t-test) and the magnitude of evoked calcium transient (dF/F0) within that subset of activated cells (0.10 ± 0.03 with PP stimulation versus 0.38 ± 0.03 without; p < 0.001; paired t-test). As a negative control, we also delivered photo-stimulation without PP activation in a subset of imaging fields, and confirmed that, as expected, no GCs were directly activated (data not shown). To test whether this rescue was specific to activation of PV-INs, we repeated the experiment with optogenetic activation of SST cells (using *Scn1a*^*+/-*^.SST-Cre mice). We found that SST activation also decreased GC responses to PP stimulation (Figure 7G-H), but to a much lesser degree (GC proportion of cells activated decreased to 0.27 ± 0.05 of baseline with PV-IN versus 0.53 ± 0.06 of baseline with SST-IN photostimulation; GC calcium transient amplitudes decreased to 0.19 ± 0.04 of baseline with PV-IN versus 0.40 ± 0.06 of baseline with SST-IN photostimulation; p < 0.01 for both comparisons, using mixed-model analyses).

## Discussion

Prior work in the field has established that Dravet syndrome (DS) is largely due to heterozygous pathogenic loss-of-function variants in *SCN1A* leading to haploinsufficiency of the voltage-gated sodium channel *α* subunit Nav1.1 (***Claes et al., 2001***). However, mechanisms by which loss of Nav1.1 leads to the epilepsy, neurodevelopmental disability and SUDEP that clinically defines DS remains under intense investigation. Work in preclinical experimental mouse models of DS (*Scn1a*^*+/-*^ mice) demonstrates temporal lobe-onset seizures (***Liautard et al., 2013***) and prominent activation of hippocampal DG during temperature-sensitive seizures (***Dutton et al., 2017***), suggesting involvement of hippocampus in seizure generation in *Scn1a*^*+/-*^ mice. In this study, we employed 2P calcium imaging in acute slice in an experimental mouse model of DS (Figure 1) to demonstrate a profound abnormality in a cortico-hippocampal circuit element strongly linked to seizures and epilepsy (Figure 2). This abnormality was not present in early postnatal mice (P14-21) yet appeared during the chronic phase of the disorder. Moreover, we found only subtle defects in PV-IN excitability (Figure 3), which did not account for the identified circuit dysfunction (Figure 4), and found no impairment in disynaptic feedforward inhibition from entorhinal cortex onto GCs (Figure 5). In contrast, we found an increased strength of monosynaptic excitatory input from entorhinal cortex to DG. Optogenetic activation of the perforant path markedly decreased the temperature threshold for seizure generation in *Scn1a*^*+/-*^ mice in vivo (Figure 6). We then targeted and corrected this circuit abnormality in vitro by recruiting inhibitory reserve via optogenetics (Figure 7). Taken together, these results demonstrate a clear locus of dysfunction despite complex circuit abnormalities and suggest unexpected plastic rearrangements in cortico-hippocampal circuity that may relate to the maintenance of chronic epilepsy and intellectual disability in DS.

### Calcium imaging enables large-scale comparison of dentate activation in wild-type and *Scn1a*^*+/-*^ mice

2P imaging allows simultaneous read-out of the activity of hundreds (and now even many thousands or more (***Stringer et al., 2019***)) of neurons, with cellular resolution and at high speeds. Our aim was to use 2P imaging for a large-scale comparison of GC activation between genotypes (*Scn1a*^*+/-*^ versus wild-type), which is a valid approach only if there is no genotype difference in the relationship between the number of action potentials that a neuron fires and its resulting calcium signal (dF/F0). This is indeed a theoretical concern as previous studies have demonstrated dysregulation of voltage gated calcium channel expression patterns within DG in rodent models as well as human patients with TLE (***Xu and Tang, 2018***). Calcium binding proteins, such as calmodulin and calbindin, may also be altered by, and play a role in, the pathological process of epilepsy (***Xu and Tang, 2018***). We therefore directly tested the ability of GCaMP7s to report action potentials in both *Scn1a*^*+/-*^ and wild-type GCs (Figure 1), confirming that we could detect single action potentials in both genotypes under our experimental conditions and that there was no difference between genotypes in the relationship between action potentials and calcium-induced fluorescence increases. That this was the case may relate to the relative homogeneity of GCs and the fact that GC cellular physiology does not appear to change in *Scn1a*^*+/-*^ mice (Table 1). We performed whole-cell patch-clamp recordings of GCs from *Scn1a*^*+/-*^ and age-matched wild-type littermate controls to confirm that intrinsic excitability was not significantly different, with no evidence of, for example, bursting in GCs in *Scn1a*^*+/-*^ mice.

### Cortico-hippocampal circuit dysfunction and ictogenesis in *Scn1a*^*+/-*^ mice

Consistent with previous literature (***Yu et al., 2013***), we found that, in wild-type mice, the response of GCs to PP stimulation is robust in early postnatal mice (P14-21), becoming sparse by the young adult timepoint (Figure 2). This developmental pattern was disrupted in *Scn1a*^*+/-*^ mice. Thus, the GC response was mildly abnormal in early postnatal mice (P14-21), whereas, relative to wild-type age-matched controls, GCs were profoundly hyper-activated in young adult mice across all conditions and analyses. Cortico-hippocampal circuit dysfunction is therefore unlikely to be the cause of epilepsy onset in early postnatal *Scn1a*^*+/-*^ mice, as spontaneous seizures in this model begin at approximately P16-18 (***Hawkins et al., 2017***), but it may contribute to ongoing seizures in the chronic phase. This inference is supported by clear evidence of temporal lobe dysfunction (***Van Poppel et al., 2012; Kasperavičiūtė et al., 2013; Gaily et al., 2013; Siegler et al., 2005; Tiefes et al., 2019***) in human patients with DS, although DS is a multifocal epilepsy and seizures do not exclusively emanate from the temporal lobe.

The mechanism underlying the developmental pattern of the cortico-hippocampal circuit deficit remains unclear. It is possible that the determinants of the cortico-hippocampal circuit dysfunction are already fully present in early postnatal *Scn1a*^*+/-*^ mice, but are masked by the robust response at this time point due to the fact that the GABAergic system remains immature (***Ben-Ari, 2002***). Alternatively, this circuit dysfunction could develop over time, as a direct consequence of the genetic lesion, an accumulated insult from ongoing seizures, a complex interaction between genotype and seizure burden (***Dutton et al., 2017; Salgueiro-Pereira et al., 2019***) or a pathologic compensation in response to loss of Nav1.1. These possibilities could be further explored using experimental manipulations to exacerbate (***Dutton et al., 2017; Hawkins et al., 2017; Salgueiro-Pereira et al., 2019***) or ameliorate (***Hawkins et al., 2017***) the epilepsy phenotype of *Scn1a*^*+/-*^ mice; subsequent testing of the excitability of the cortico-hippocampal circuit could help parse the contribution of the genotype (common across cohorts) versus, for example, the effect of ongoing seizures, to disease severity. Conversely, it would be of interest to focally rescue *Scn1a* levels exclusively in dentate gyrus and/or entorhinal cortex of *Scn1a*^*+/-*^ mice (***Colasante et al., 2020***), to test whether the cortico-hippocampal circuit would remain abnormal (assuming continued seizures involving the hippocampus).

The failure of Hm1a to rescue this circuit pathology (Figure 4) highlights the conclusion that, by the young adult time point, the mechanism is more complex than Nav1.1 insufficiency (although it is the case that Hm1a application is not equivalent to doubling Na+ current density). This distinction has treatment implications: recent preclinical work using antisense oligonucleotide (ASOs) to increase *Scn1a* expression showed promising improvement in the disease phenotype when delivered at P2 or P14, prior to seizure onset (***Han et al., 2020***). However, it remains to be seen whether boosting Nav1.1 expression levels with ASOs would be effective in the chronic phase of the disorder, after secondary circuit deficits may already be in place. Another recent study (***Yamagata et al., 2020***), using CRISPR/dCas9-based gene activation to increase Nav1.1 expression in GABAergic interneurons at P28, found an increased temperature threshold of seizure induction; however, the treated mice appeared to have a marked increase in early mortality (prior to intervention) compared to the comparison groups, rendering this finding difficult to interpret.

A central mechanistic hypothesis in acquired temporal lobe epilepsy (TLE) is that epileptogenesis results from uncontrolled excitatory input to vulnerable downstream hippocampal areas, leading to damage to CA3 and seizure propagation through the hippocampal network (***Dengler et al., 2017; Behr et al., 1998***). In chronic acquired TLE after brain injury (such as due to status epilepticus induced by a chemoconvulsant agent) in rodents, the current operating model is a transient (weeks) impairment of cortico-hippocampal circuit function during the so-called latent period, followed by reconstitution of normal/near-normal operations, then secondary temporoammonic (TA) pathway failure and/or chronic impairment of DG activity (***Dengler et al., 2017; Ang et al., 2006***). Our findings suggest that impaired DG function may be a convergent circuit-level feature shared across acquired TLE and DS in the chronic phase, although the inciting insult is completely different between these epilepsy types. We did not directly investigate the TA pathway in this study, although our in vivo optogenetic stimulation of cell bodies within entorhinal cortex (Figure 6) likely activated TA as well as PP projections to hippocampus. Disambiguating the relative contribution of PP and TA pathway to ictogenesis in *Scn1a*^*+/-*^ mice would be an interesting future direction.

### Excess synaptic excitation onto dentate gyrus granule cells

As PV-INs in DG have been shown to powerfully regulate the GC response to entorhinal cortical input (***Lee et al., 2016***) and GABAergic interneurons are known to be abnormal in *Scn1a*^*+/-*^ mice (***Mistry et al., 2014; Bechi et al., 2012; Yu et al., 2006; Ogiwara et al., 2013; Tsai et al., 2015***), we initially hypothesized that PV-IN dysfunction would be the primary mechanism underlying the GC hyperactivation observed here (Figure 2). However, our analysis of PV-INs in DG showed only a relatively subtle impairment in action potential generation in young adult mice, characterized by run-down of action potential peak during prolonged trains at high frequency in response to large-amplitude current injections, likely due to accumulation of sodium channel inactivation (Figure 3). The physiological relevance of these abnormalities are unclear and are likely insufficient to explain the profound circuit-level breakdown in dentate function that we observed, as PV-INs in *Scn1a*^*+/-*^ mice were still able to fire at >200 Hz for trains of 10+ action potentials. This inference was supported by the more critical observations that circuit dysfunction persisted in the presence of saturating concentrations of PTX combined with the finding that the amplitude of the evoked disynaptic feedforward IPSC onto GCs was the same between genotypes (or was perhaps even larger in *Scn1a*^*+/-*^ mice; Figure 5). Previous studies have shown impaired firing properties of hippocampal PV-INs, although these were in younger (:S P21) mice (***Tsai et al., 2015; Hedrich et al., 2014***).Near-normalization of high frequency firing of DG PV-INs in young adult *Scn1a*^*+/-*^ mice is consistent with prior work demonstrating that PV-IN dysfunction (in layer 2/3 barrel cortex) is limited to early developmental time points (***Favero et al., 2018***). Our finding is also consistent with recent work demonstrating that sensorimotor cortex PV-INs in adult *Scn1a*^*+/-*^ mice exhibit normal activity levels during quiet wakefulness in vivo (***Tran et al., 2020***).

Our measurement of evoked excitatory and disynaptic inhibitory post-synaptic currents onto GCs also points away from the sole locus of circuit dysfunction in chronic DS being impaired inhibition, and instead suggests that the underlying mechanism of *Scn1a*^*+/-*^ GC hyperactivation is more complex, and may involve excessive synaptic excitation onto those GCs (Figure 5A-E), the basis of which is unclear. This result is also consistent with our observation of durable hyperactivation of GCs in *Scn1a*^*+/-*^ mice relative to wild-type even in the presence of GABA_A_ receptor blockade (using PTX), which also points to excess excitation as opposed to impaired inhibition (Figure 5F). Increased synaptic excitation onto GCs in *Scn1a*^*+/-*^ mice could arise from presynaptic and/or post-synaptic mechanisms. Nav1.1 is likely expressed within a subset of dentate-projecting entorhinal cortical cells (***Ogiwara et al., 2013***), and it is possible that these cells are rendered hyperexcitable due to a similar compensatory mechanism that underlies normalization of neocortical (and DG) PV-INs. Our finding is consistent with previous studies showing hyperexcitability and increased synaptic excitation in downstream hippocampal CA1 (***Liautard et al., 2013; Hedrich et al., 2014; Han et al., 2012; Gu et al., 2014***). In contrast, in a prior study showing increased evoked GC activation following status epilepticus-induced TLE, dentate hyperactivation in the epilepsy condition relative to control was eliminated by PTX (***Dengler et al., 2017***). This illustrates how disparate underlying epilepsy types (genetic mutation versus brain injury) could converge at the level of the circuit (DG hyperactivity) via disparate mechanisms (impaired inhibition versus excessive excitation).

### Rescue of cortico-hippocampal circuit excitability via targeted recruitment of PVINs

To test the impact of local inhibitory interneurons on the recruitment of GCs by PP input, we combined 2P imaging with optogenetic stimulation with the red-shifted channelrhodopsin variant ChrimsonR (***Klapoetke et al., 2014***) (Figure 7). This technique enabled us to perform simultaneous imaging (of anatomically-defined GCs) and photostimulation (of genetically-defined interneuron subtypes; i.e. PV-INs or SST-INs) within the same slice. We found that targeted PV-IN activation in acute slice significantly decreased GC activation in response to PP input (Figure 7). Our photostimulation likely activated not only PV-IN cell bodies but also ChrimsonR-expressing fibers. Previous work has shown anatomic differences in the axon initial segment within *Scn1a*^*+/-*^ PV-INs (***Favero et al., 2018***), with implications on axonal function that are not yet clear. Photoactivation of distal fibers could mask a proximal deficit in these cells that could pose an obstacle to a theoretical future soma-targeted therapy.

Optogenetic activation of dentate PV-INs has been shown to truncate seizures in vivo in a mouse model of TLE (***Krook-Magnuson et al., 2013***). It remains to be seen whether a similar strategy may be successfully employed against temporal lobe-onset (or other) seizures in *Scn1a*^*+/-*^ mice. Our data shows that, while DG PV-INs do exhibit minor abnormalities in spike generation during the chronic phase of the disorder, these cells remain capable of discharging short trains of action potentials at high frequency (> 200 Hz) such that there is sufficient inhibitory “reserve” that would enable these cells to be exogenously recruited to balance aberrant excitatory inputs.

Improved understanding of the mechanisms of ictogenesis in DS is critical towards the development of novel therapies for what is currently an incurable and treatment-resistant disorder associated with high mortality. Our results indicate complex network dysfunction underlying abnormalities in a node that has been strongly implicated in epilepsy and that cannot be clearly attributed solely to dysfunction of GABAergic subsystems. We provide evidence to suggest convergent cellular and network mechanisms underlying temporal lobe seizures across diverse forms of epilepsy that could potentially be targeted for seizure suppression or interruption in DS. As sparse GC firing is also thought to be crucial for cognitive functions of the hippocampus, such as pattern separation and memory encoding (***Leutgeb et al., 2007; Treves et al., 2008; Rolls, 2010; McHugh et al., 2007***), correction of this circuit dysfunction may also potentially modulate the severe and chronic intellectual disability characteristic of DS which is even more difficult to approach than epilepsy.

## Methods and Materials

All experiments were carried out according to protocols approved by the Institutional Animal Care and Use Committee at the Children’s Hospital of Philadelphia, in accordance with ethical guidelines set forth by the National Institutes of Health.

### Experimental animals

Male and female mice were used in equal proportion. Mice were weaned at P21 and were subsequently group-housed to the extent possible in cages containing up to five mice. They were maintained on a 12 hour light/dark cycle with access to food and water ad libitum. All mice were genotyped via PCR analysis of a tail snip obtained at P7.

The *Scn1a*^*+/-*^ mice used in this study have a targeted deletion of the first exon of the *Scn1a* gene, with a resulting null allele and 50% decrease in Nav1.1 protein (***Mistry et al., 2014; Miller et al., 2014***). To generate mice for experimental use, male *Scn1a*^*+/-*^ mice on a 129S6.SvEvTac background (RRID: MMRRC_037107-JAX) were crossed to female mice on a C57BL6/J background, either wild-type C57BL6/J mice (RRID: IMSR_JAX:000664) or Cre-driver lines (see below). The resulting progeny have a mixed 50:50 129S6:BL6/J background, on which the DS phenotype has been extensively characterized (***Mistry et al., 2014; Hawkins et al., 2017; Han et al., 2020; Miller et al., 2014***).

For experiments requiring targeted viral expression in, or visualization of, PV-INs, female PV-Cre.tdT double-heterozygous mice were generated from a cross between homozygous tdTomato reporter/Ai14 mice (Rosa-CAG-LSL-tdTomato; RRID: IMSR_JAX:007914) and homozygous PV-Cre mice (B6;129P2-Pvalbtm1(cre)Arbr; RRID: IMSR_JAX:008069). These mice are on a C57BL6/J background. The F1 progeny include *Scn1a*^*+/-*^.PV-Cre.tdT and PV-Cre.tdT mice (1:8 predicted Mendelian ratio for each). These mice express tdT in PV cells, which facilitates targeted experiments. A similar strategy was used for experiments targeting viral expression to SST-INs, with SST-Cre mice (B6J.Cg-Ssttm2.1(cre)Zjh; RRID: IMSR_JAX:028864).

### Viral injections

Subdural viral injections (early postnatal mice): To achieve GCaMP expression in early postnatal mice, subdural viral injections68 were performed on mice at age P0-2. Mice were anesthetized by cooling on ice until cessation of motor activity. The scalp was sterilized using EtOH. Virus was injected halfway between lambda and bregma and halfway between the midline and the eye, using a 10 µL syringe (Hamilton) and a 33G beveled needle (World Precision Instruments). Injections of GCaMP (AAV9.syn.GCaMP7s.WPRE; Addgene) were performed bilaterally using 1 µL of virus at a genomic titer of 1.5 × 1013 cfu/mL.

Stereotaxic viral injections (young adult mice): Mice were anesthetized by inhalation of isoflurane (4% induction, 1-2% maintenance) in oxygen. Anesthesia depth was monitored by response to toe pinch and by breathing rate. Animals were injected subcutaneously with buprenorphine (0.5-1.0 mg/kg, sub-Q), cefazolin (500 mg/kg, sub-Q), and meloxicam (5 mg/kg, sub-Q), for post-operative analgesia and anti-bacterial prophylaxis. Mice were placed in a stereotaxic apparatus (Kopf Instruments) while resting on a heating pad. After removal of fur and sterilization of the scalp, a scalpel was used to expose the skull and target regions were identified in relation to skull surface landmarks (lambda and bregma). Craniotomies were made using a hand-held drill. Injections were performed with 200-250 nL of virus per site, at a rate of 100 nL/minute, using a 10 µL syringe (Hamilton) and a 33G beveled needle (World Precision Instruments). All injections were unilateral (on the right hemisphere). Injection coordinates for entorhinal cortex (relative to Bregma, in mm) were -4.6 (A/P); 3.0 (M/L); 4.1 and 4.6 (D/V). Injections in dorsal and ventral DG were achieved 10-12 injections per animal, spanning from -1.5 to -3.8 A/P. The viruses and genomic titers were as follows: AAV9.syn.GCaMP7s.WPRE (Addgene), 3 × 1012 cfu/mL; AAV9.syn.FLEX.ChrimsonR.tdT (Addgene), 2.3 × 1012 cfu/mL; AAV9.CamKIIa.hChR2(H134R)-eYFP.WPRE.hGH (Addgene), 1.8 × 1013 cfu/mL. Experiments were performed at least 3 weeks after injection (to allow for virus expression).

### Acute slice preparation

Mice were deeply anesthetized using inhaled isoflurane. Adult mice were perfused transcardially with 5 mL ice-cold sucrose-based cutting solution. Brains were removed and immediately transferred to ice-cold sucrose solution (in mM: NaCl, 87; sucrose, 75; KCl, 2.5; CaCl2, 1.0; MgSO4, 2.0; NaHCO3, 26; NaH2PO4, 1.25; glucose, 10), and equilibrated with 95% O2 and 5% CO2. Hippocampalentorhinal cortex (HEC) slices were obtained to preserve within the slice the PP connection between the entorhinal cortex and the hippocampus (***Xiong et al., 2017***). A mid-sagittal cut was made to divide the hemispheres, and one hemisphere was mounted onto an angled agar block. The blade therefore passed through the block at an acute (15°) downward angle from rostral to caudal, resulting in modified horizontal/axial HEC slices. Slices were sectioned at a 300 µm thickness using a Leica VT-1200S vibratome. Slices were transferred to a holding chamber containing the same sucrose solution. After a recovery period of 30 minutes at 30-32°C, the holding chamber was removed from the water bath and allowed to cool to room temperature. For data acquisition, slices were transferred to a recording chamber (shielded from light in the case of opsin-containing slices) and continuously perfused with oxygenated artificial cerebrospinal fluid (ACSF; in mM: NaCl, 125; KCl, 2.5; CaCl2, 2.0; MgSO4, 1.0; NaHCO3, 26; NaH2PO4, 1.25; glucose, 10), continuously bubbled with 95% O2 and 5% CO2, at a rate of approximately 3 mL/min, heated to 30-32 °C.

### Perforant path stimulation

The perforant path was stimulated using a 125 µm diameter concentric bipolar microelectrode (FHC Inc. NC0950490). The electrode was placed approximately 100 µm from the hippocampal fissure, on the side of the entorhinal cortex, adjacent to the region between the suprapyramidal blade and the apex of the dentate gyrus granule cell layer, so as not to directly stimulate GCs. Since GCs are known to be more readily activated by PP stimulation in younger mice (***Yu et al., 2013***), we selected different ranges of stimulation intensity for the two groups (25-400 µA for early postnatal; 100-500 µA for young adult).

### Whole cell slice recordings

DG GCs were identified by localization within the GCL and morphology confirmed by electrophysiological discharge pattern. PV-INs were identified by endogenous tdT expression visualized with epifluorescence and were typically large cells located at the border between the granule cell layer and the hilus. Whole cell voltage- and current-clamp recordings were obtained using borosilicate glass electrodes, pulled using a P-97 puller (Sutter Instruments) for a tip resistance of 3-4 MΩ for voltage clamp and 4-6 MΩ for current clamp.

For PV-IN patching, the pipette solution was a K-Gluconate solution that contained, in mM: K-gluconate, 130; KCl, 6.3; EGTA, 0.5; MgCl2, 1.0; HEPES, 10; Mg-ATP, 4.0; Na-GTP, 0.3; pH adjusted to 7.30 with KOH and osmolarity adjusted to 285 mOsm with 30% sucrose. For characterization of intrinsic electrophysiological properties of GCs, the pipette solution was a chloride-free solution that contained, in mM: K-gluconate, 150; MgCl2, 1.0; HEPES, 10; Mg-ATP, 5.0; Na-GTP, 0.3; pH adjusted to 7.40 with KOH and osmolarity adjusted to 285 mOsm with 30% sucrose. To measure both EPSCs and IPSCs in GCs, whole cell recordings were obtained in an independent set of GCs with a cesium-based pipette solution that contained, in mM: Cs Methylsulfonate, 125; HEPES, 15; EGTA 0.5; Mg-ATP, 2.0; Na-GTP, 0.3; phosphocreatine-Tris2, 10; QX 314, 2.0; TEA, 2; pH adjusted to 7.35 with CsOH and osmolarity adjusted to 295 mOsm with 30% sucrose.

Signals were sampled at 50 kHz, amplified with a MultiClamp 700B amplifier (Molecular Devices), filtered at 10 kHz, digitized using a DigiData 1550, and acquired using pClamp10 software. Data were not included for final analysis if the cell had a membrane potential greater (less negative) than -60 mV (for GCs) or greater than -50 mV (for PV-INs), or if access resistance was greater than 30 MΩ. Series resistance compensation (bridge balance) was applied throughout current clamp experiments with readjustments as necessary. Reported values for membrane potential and AP threshold are not corrected for the liquid junction potential.

### Cell-attached slice recordings

Quantum Dots (***Andrásfalvy et al., 2014***) were employed to enable visualization of the pipette tip via two-photon imaging. Dried Quantum Dots (625 nm emission) were reconstituted in hexane. Freshly pulled borosilicate glass electrodes (7-9 MΩ) were dipped 5-10 times into the solution, while maintaining positive pressure to avoid clogging the pipette tip. These cell-attached recordings were performed with 100 µM PTX added to the bath solution to increase the proportion of cells responsive to PP stimulation.

### 2P calcium imaging and combined optogenetic stimulation in acute slice

Imaging was performed using a customized two-photon laser scanning microscope (Bruker Ultima) equipped with a resonant scanner (Cambridge Technologies) and a MaiTai DeepSee Ti:Sapphire mode-locked pulsed infrared laser (SpectraPhysics). GCaMP7s was imaged at 920 nm with a gallium arsenide phosphide (GaAsP) photodetector (H7422-40; Hamamatsu) through a 25X/0.95-NA water immersion objective (Leica). Laser power at the sample was estimated using a photodiode power sensor (Thorlabs, S120C). Laser power typically ranged from 3-5 mW (at the pickoff); for imaging in the acute brain slice preparation. For each stimulation, 500 frames were collected with a frame period of 33.8 ms (30 Hz) and a resolution of 512 × 512 pixels.

Full-field photostimulation was delivered through the objective lens with a high-powered red (660 nm) LED (M660L4; Thorlabs). This was mounted to the epifluorescence port of the microscope and routed to the sample below the PMTs using a custom notched dichromic mirror/polychromic beam splitter with reflectance from 660 ± 20 nm (ZT660/40; Chroma). LEDs were controlled via an LED driver (LEDD1B; Thorlabs) driven by a TTL pulse from and synchronized with either the imaging acquisition software (PrairieView) or electrophysiology data acquisition software (pClamp). Irradiance at the specimen was measured through the imaging objective, with measurements correspond to defined values of current applied by the LED driver. A light power density of 26 mW/mm2 was used for all experiments.

### Slice pharmacology

Picrotoxin (Tocris) was used at a saturating concentration of 100 µM in the ACSF solution. Hm1a (Alomone Labs STH-601) was used at a 50 nM final concentration in 0.025% BSA. Both drugs were perfused in at 3 mL/min after a baseline recording was obtained. Wash-on times were 10 minutes for PTX and 10-25 minutes for Hm1a.

### Immunohistochemistry

Mice were deeply anesthetized with isoflurane and transcardially perfused with ice-cold PBS followed by 4% paraformaldehyde (PFA). Brains were removed and post-fixed in PFA overnight, and equilibrated in a 30% sucrose solution. We then cut 40 µm sections on a frozen sliding microtome (American Optical). After washing, slices were stained with DAPI (Fisher Scientific), cover-slipped, and imaged using an upright epi-fluorescence microscope (Nikon Instruments).

### In vivo optogenetics and hyperthermic seizure generation

Following virus injection, mice were implanted with a fiberoptic cannula (200 µm core diameter, 240 µm outer diameter, 0.22 NA, flat tip) coupled to a Zirconia ferrule (1.25 mm outer diameter; Doric Lenses). This was implanted above the entorhinal cortex (unilateral, on the right hemisphere). Implant coordinates (relative to Bregma) were -4.6 (A/P); 3.0 (M/L); 3.8 (D/V). The implant was secured to the skull with a layer of adhesive cement (CB metabond) followed by dental cement (Patterson dental) and then capped prior to use.

In vivo optogenetic stimulation was performed approximately 3 weeks after surgery to allow for virus expression and recovery from surgery. Blue light was generated by a 473 nm laser (Shanghai Laser Optics Century Co.) and delivered to mice through a fiberoptic patch cord (0.22 NA, 200 µm core diameter; Doric Lenses) coupled to the implanted fiberoptic cannula via a connecting plastic sleeve (Precision Fiber Products). Blue laser output was controlled using a pulse generated (Master-8) to deliver 5-ms light pulse trains at a rate of 20 Hz, 5 seconds on / 5 seconds off. Light power was initially 5 mW (39.8 mW/mm2 at the tip of the fiberoptic); if no seizure was elicited after 5 minutes the power was increased to 10 mW (79.6 mW/mm2 at the tip of the fiberoptic).

Seizures were elicited by passive elevation of core body termperature. Temperature was monitored continuously via rectal probe (Physitemp), which was secured in place so as to facilitate stable recordings during freely moving behavior in the recording chamber. Mice were placed in a plexiglass chamber at room temperature and monitored for 5 minutes to establish a stable baseline. Mice were then transferred to a second chamber that was pre-warmed with an temperature controller (Physitemp TCAT-2AC). The recording was continued either until a behavioral seizure was observed, or until the body temperature reached 42.5 °C, upon which the mouse was immediately removed from the chamber and cooled using ice.

### Analysis of whole-cell electrophysiology data

All analysis was performed blind to genotype using Matlab (Mathworks) or Python (custom software using Python 3.7 and the pyABF model) with quality control using manual confirmation in Clampfit (pCLAMP). Resting membrane potential (Vm) was calculated using the average value of a 1 s sweep with no direct current injection. Input resistance (Rm) was calculated using the average response to a small hyperpolarizing step near rest, using Rm = V/I for each sweep. Spike height refers to the absolute maximum voltage value of an individual AP. For GCs, spikes were defined as having at least a 40 mV amplitude (difference between spike height and AP threshold), and a spike height overshooting 0 mV. For PV-INs, spikes were defined as having a clear threshold, at which the derivative of the voltage (dV/dt) is greater than 10 mV/ms, and a spike height overshooting 0 mV. AP threshold was calculated as the value at which the derivative of the voltage (dV/dt) first reached 10 mV/ms. Rheobase was determined as the minimum current injection that elicited APs using a 600 ms sweep at 10-50 pA intervals. AP half-width is defined as the width of the AP (in ms) at half-maximal amplitude (half the voltage difference between the AP threshold and peak). Maximal instantaneous firing was calculated as the inverse of the smallest interspike interval elicited during a suprathreshold 600 ms current injection. Maximal steady-state firing was defined as the maximal mean firing frequency during a suprathreshold 600 ms current injection. All quantification of single spike properties was done using the first AP elicited at rheobase (for GCs) or at twice rheobase (for PV-INs). I-f plots were created using the steady-state firing calculated for each current step, counting failures as 0 for subsequent current steps.

### Analysis of 2P imaging data

All conditions under each field of view were grouped together as a raw data file. Motion was minimal or absent in most cases, but there was some drift during longer imaging sessions, such as with pharmacologic application. Therefore, within each raw data file, all individual frames were motion corrected using the Non-Rigid Motion Correction (NoRMCorre) Matlab toolbox (***Pnevmatikakis, 2019***). The first 200 frames were used for the template. The motion corrected individual frames were averaged to obtain an average image for the respective field of view. Regions of interest (ROIs) were identified from the average image. For the Hm1a imaging data, which required keeping the slice in plane for a prolonged wash-in, ROIs encompassed the entire granule cell layer, the boundaries of which were manually drawn. For all other data, ROIs were individual GCs, and were identified using the NeuroSeg70, a Matlab toolbox with both automatic and manual options for selection of regions of cells; we primarily relied upon the manual option as we observed that the automated algorithm was not able to effectively identify the densely packed GCs.

The raw traces from each identified cell were extracted and low pass filtered. The first 50 frames of each trace in each condition were considered as a baseline to compute dF/F0 for each trace. Peaks of the traces were detected using the findpeaks function in Matlab. A threshold of (mean+3*standard deviation) was used to define significant peaks. A cell was detected as active if a significant peak is shown in the respective trace for that particular condition. The results were summarized for each condition by both the proportion of active cells and the individual significant peak amplitudes.

A mixed modeling approach was implemented for statistical analysis of data across populations. This was performed due to the hierarchical experimental design, in which sources of variance could arise from effects of imaging fields, slices, and mice. The proportion of activated cells were modeled as a binomial distribution with a probit link function. The amplitude of peaks were modeled as a normal distribution with a logarithmic link function.

For spike deconvolution, spikes were extracted using constrained non-negative matrix factorization (***Pnevmatikakis et al., 2016***). The deconvolution parameters such as rise time and decay time were fine-tuned based on the ground truth data. A cell was defined as active if there were one or more spikes detected. Deconvolution yielded a single spike or multiple spikes based on the perforant path stimulation pattern (1 pulse versus 4 pulses). Irrespective of the stimulation, the total deconvolved spikes are derived by summing the individual deconvolved spikes obtained during the total time range of stimulation.

For other analyses involving comparisons within a given imaging field (e.g. response to PP stimulation alone versus PP stimulation plus optogenetic activation of PV-INs), paired t-tests were performed when comparing two conditions. For comparing more than two conditions, a one-way ANOVA was performed followed by post hoc paired t-tests with Bonferroni correction. Statistical significance was defined as p < 0.05 with p values reported exactly.

## Acknowledgments

This work was supported by NIH NINDS Research Education Grant (R25) to J.M., and NIH NINDS K08 NS097633, NIH NINDS R01 NS110869, the Dana Foundation David Mahoney Neuroimaging Program research grant, and the Burroughs Wellcome Fund Career Award for Medical Scientists to E.M.G. We would also like to thank the Women’s Committee of The Children’s Hospital of Philadelphia for support. Statistical support was obtained from the Center for Human Phenomic Science (CHPS; CTSA grant UL1TR001878). We thank the GENIE project and the Janelia Research Campus of the HHMI for distribution of GCaMP7 and Ed Boyden at MIT for distribution of ChrimsonR

## References

Amaral DG, Scharfman HE, Lavenex P. The dentate gyrus: fundamental neuroanatomical organization (dentate gyrus for dummies). In: Progress in Brain Research, vol. 163 Elsevier; 2007.p. 3–790. https://linkinghub.elsevier.com/retrieve/pii/S0079612307630015, dOI: 10.1016/S0079-6123(07)63001-5.

Andrásfalvy BK, Galiñanes GL, Huber D, Barbic M, Macklin JJ, Susumu K, Delehanty JB, Huston AL, Makara JK, Medintz IL. Quantum dot-based multiphoton fluorescent pipettes for targeted neuronal electrophysiology. Nature methods. 2014 12; 11(12):1237–1241. doi: 10.1038/nmeth.3146, pMID: 25326662 PMCID: PMC4245189.

Ang CW, Carlson GC, Coulter DA. Massive and Specific Dysregulation of Direct Cortical Input to the Hippocampus in Temporal Lobe Epilepsy. Journal of Neuroscience. 2006 11; 26(46):11850–11856. doi: 10.1523/JNEUROSCI.2354-06.2006.

Bechi G, Scalmani P, Schiavon E, Rusconi R, Franceschetti S, Mantegazza M. Pure haploinsufficiency for Dravet syndrome NaV1.1 (SCN1A) sodium channel truncating mutations. Epilepsia. 2012; 53(1):87–100. doi: 10.1111/j.1528-1167.2011.03346.x.

Behr J, Lyson KJ, Mody I. Enhanced Propagation of Epileptiform Activity Through the Kindled Dentate Gyrus. Journal of Neurophysiology. 1998 4; 79(4):1726–1732. doi: 10.1152/jn.1998.79.4.1726, publisher: American Physiological Society.

Ben-Ari Y. Excitatory actions of gaba during development: the nature of the nurture. Nature Reviews Neuroscience. 2002 9; 3(9):728–739. doi: 10.1038/nrn920, number: 9 publisher: Nature Publishing Group.

Chawla MK, Guzowski JF, Ramirez-Amaya V, Lipa P, Hoffman KL, Marriott LK, Worley PF, McNaughton BL, Barnes CA. Sparse, environmentally selective expression of Arc RNA in the upper blade of the rodent fascia dentata by brief spatial experience. Hippocampus. 2005; 15(5):579–586. doi: 10.1002/hipo.20091, pMID: 15920719.

Cheah CS, Yu FH, Westenbroek RE, Kalume FK, Oakley JC, Potter GB, Rubenstein JL, Catterall WA. Specific deletion of NaV1.1 sodium channels in inhibitory interneurons causes seizures and premature death in a mouse model of Dravet syndrome. Proceedings of the National Academy of Sciences. 2012 9; 109(36):14646– 14651. doi: 10.1073/pnas.1211591109.

Claes L, Del-Favero J, Ceulemans B, Lagae L, Van Broeckhoven C, De Jonghe P. De novo mutations in the sodium-channel gene SCN1A cause severe myoclonic epilepsy of infancy. American Journal of Human Genetics. 2001 6; 68(6):1327–1332. doi: 10.1086/320609, pMID: 11359211 PMCID: PMC1226119.

Colasante G, Lignani G, Brusco S, Di Berardino C, Carpenter J, Giannelli S, Valassina N, Bido S, Ricci R, Castoldi V, Marenna S, Church T, Massimino L, Morabito G, Benfenati F, Schorge S, Leocani L, Kullmann DM, Broccoli V. dCas9-Based Scn1a Gene Activation Restores Inhibitory Interneuron Excitability and Attenuates Seizures in Dravet Syndrome Mice. Molecular Therapy: The Journal of the American Society of Gene Therapy. 2020 1; 28(1):235–253. doi: 10.1016/j.ymthe.2019.08.018, pMID: 31607539 PMCID: PMC6952031.

Dana H, Sun Y, Mohar B, Hulse BK, Kerlin AM, Hasseman JP, Tsegaye G, Tsang A, Wong A, Patel R, Macklin JJ, Chen Y, Konnerth A, Jayaraman V, Looger LL, Schreiter ER, Svoboda K, Kim DS. High-performance calcium sensors for imaging activity in neuronal populations and microcompartments. Nature Methods. 2019 7; 16(7):649–657. doi: 10.1038/s41592-019-0435-6.

De Stasi AM, Farisello P, Marcon I, Cavallari S, Forli A, Vecchia D, Losi G, Mantegazza M, Panzeri S, Carmignoto G, Bacci A, Fellin T. Unaltered Network Activity and Interneuronal Firing During Spontaneous Cortical Dynamics In Vivo in a Mouse Model of Severe Myoclonic Epilepsy of Infancy. Cerebral Cortex. 2016 4; 26(4):1778–1794. doi: 10.1093/cercor/bhw002.

Dengler CG, Yue C, Takano H, Coulter DA. Massively augmented hippocampal dentate granule cell activation accompanies epilepsy development. Scientific Reports. 2017 2; 7:42090. doi: 10.1038/srep42090.

Diamantaki M, Frey M, Berens P, Preston-Ferrer P, Burgalossi A. Sparse activity of identified dentate granule cells during spatial exploration. eLife. 2016; 5. https://www.ncbi.nlm.nih.gov/pmc/articles/PMC5077296/, doi: 10.7554/eLife.20252, pMID: 27692065 PMCID: PMC5077296.

Dutton SB, Makinson CD, Papale LA, Shankar A, Balakrishnan B, Nakazawa K, Escayg A. Preferential inactivation of Scn1a in parvalbumin interneurons increases seizure susceptibility. Neurobiology of Disease. 2013 1; 49:211–220. doi: 10.1016/j.nbd.2012.08.012.

Dutton SBB, Dutt K, Papale LA, Helmers S, Goldin AL, Escayg A. Early-life febrile seizures worsen adult phenotypes in Scn1a mutants. Experimental Neurology. 2017 7; 293:159–171. doi: 10.1016/j.expneurol.2017.03.026.

Erum JV, Dam DV, Deyn PPD. PTZ-induced seizures in mice require a revised Racine scale. Epilepsy Behavior. 2019 6; 95:51–55. doi: 10.1016/j.yebeh.2019.02.029, pMID: 31026782.

Ewell LA, Jones MV. Frequency-Tuned Distribution of Inhibition in the Dentate Gyrus. Journal of Neuroscience. 2010 9; 30(38):12597–12607. doi: 10.1523/JNEUROSCI.1854-10.2010.

Favero M, Sotuyo NP, Lopez E, Kearney JA, Goldberg EM. A Transient Developmental Window of Fast-Spiking Interneuron Dysfunction in a Mouse Model of Dravet Syndrome. The Journal of Neuroscience. 2018 9; 38(36):7912–7927. doi: 10.1523/JNEUROSCI.0193-18.2018.

Gaily E, Anttonen AK, Valanne L, Liukkonen E, Träskelin AL, Polvi A, Lommi M, Muona M, Eriksson K, Lehesjoki AE. Dravet syndrome: new potential genetic modifiers, imaging abnormalities, and ictal findings. Epilepsia. 2013 9; 54(9):1577–1585. doi: 10.1111/epi.12256, pMID: 23808377.

van Gassen KLI, Hessel EVS, Ramakers GMJ, Notenboom RGE, Wolterink-Donselaar IG, Brakkee JH, Godschalk TC, Qiao X, Spruijt BM, van Nieuwenhuizen O, de Graan PNE. Characterization of febrile seizures and febrile seizure susceptibility in mouse inbred strains. Genes, Brain, and Behavior. 2008 7; 7(5):578–586. doi: 10.1111/j.1601-183X.2008.00393.x, pMID: 18363854.

Goff KM, Goldberg EM. Vasoactive intestinal peptide-expressing interneurons are impaired in a mouse model of Dravet syndrome. eLife. 2019; 8. https://www.ncbi.nlm.nih.gov/pmc/articles/PMC6629374/, doi: 10.7554/eLife.46846, pMID: 31282864 PMCID: PMC6629374.

Gu F, Hazra A, Aulakh A, Žiburkus J. Purinergic control of hippocampal circuit hyperexcitability in Dravet syndrome. Epilepsia. 2014; 55(2):245–255. doi: 10.1111/epi.12487.

Han S, Tai C, Westenbroek RE, Yu FH, Cheah CS, Potter GB, Rubenstein JL, Scheuer T, de la Iglesia HO, Catterall WA. Autistic-like behaviour in Scn1a+/ mice and rescue by enhanced GABA-mediated neurotransmission. Nature. 2012 9; 489(7416):385–390. doi: 10.1038/nature11356.

Han Z, Chen C, Christiansen A, Ji S, Lin Q, Anumonwo C, Liu C, Leiser SC, Meena Aznarez I, Liau G, Isom LL. Antisense oligonucleotides increase Scn1a expression and reduce seizures and SUDEP incidence in a mouse model of Dravet syndrome. Science Translational Medicine. 2020 8; 12u558). https://stm.sciencemag.org/content/12/558/eaaz6100, doi: 10.1126/scitranslmed.aaz6100, publisher: American Association for the Advancement of Science section: Research Article PMID: 32848094.

Hawkins NA, Anderson LL, Gertler TS, Laux L, George AL, Kearney JA. Screening of conventional anticonvulsants in a genetic mouse model of epilepsy. Annals of Clinical and Translational Neurology. 2017 4; 4(5):326–339. doi: 10.1002/acn3.413, pMID: 28491900 PMCID: PMC5420810.

Hedrich UBS, Liautard C, Kirschenbaum D, Pofahl M, Lavigne J, Liu Y, Theiss S, Slotta J, Escayg A, Dihné M, Beck H, Mantegazza M, Lerche H. Impaired Action Potential Initiation in GABAergic Interneurons Causes Hyperexcitable Networks in an Epileptic Mouse Model Carrying a Human NaV1.1 Mutation. Journal of Neuroscience. 2014 11; 34(45):14874–14889. doi: 10.1523/JNEUROSCI.0721-14.2014, publisher: Society for Neuroscience section: Articles PMID: 25378155.

Houser CR. Interneurons of the dentate gyrus: an overview of cell types, terminal fields and neurochemical identity. In: Scharfman HE, editor., Progress in Brain Research, vol. 163 of The Dentate Gyrus: A Comprehensive Guide to Structure, Function, and Clinical Implications Elsevier; 2007.p. 217–811. http://www.sciencedirect.com/science/article/pii/S0079612307630131, dOI: 10.1016/S0079-6123(07)63013-1.

Jiao J, Yang Y, Shi Y, Chen J, Gao R, Fan Y, Yao H, Liao W, Sun XF, Gao S. Modeling Dravet syndrome using induced pluripotent stem cells (iPSCs) and directly converted neurons. Human Molecular Genetics. 2013 11; 22(21):4241–4252. doi: 10.1093/hmg/ddt275.

Kasperavičiūte? D, Catarino CB, Matarin M, Leu C, Novy J, Tostevin A, Leal B, Hessel EVS, Hallmann K, Hildebrand MS, Dahl HHM, Ryten M, Trabzuni D, Ramasamy A, Alhusaini S, Doherty CP, Dorn T, Hansen J, Krämer G, Steinhoff BJ, et al. Epilepsy, hippocampal sclerosis and febrile seizures linked by common genetic variation around SCN1A. Brain. 2013 10; 136(10):3140–3150. doi: 10.1093/brain/awt233.

Klapoetke NC, Murata Y, Kim SS, Pulver SR, Birdsey-Benson A, Cho YK, Morimoto TK, Chuong AS, Carpenter EJ, Tian Z, Wang J, Xie Y, Yan Z, Zhang Y, Chow BY, Surek B, Melkonian M, Jayaraman V, Constantine-Paton M, Wong GKS, et al. Independent optical excitation of distinct neural populations. Nature Methods. 2014 3; 11(3):338–346. doi: 10.1038/nmeth.2836.

Krook-Magnuson E, Armstrong C, Oijala M, Soltesz I. On-demand optogenetic control of spontaneous seizures in temporal lobe epilepsy. Nature Communications. 2013 6; 4(1):1376. doi: 10.1038/ncomms2376.

Lee CT, Kao MH, Hou WH, Wei YT, Chen CL, Lien CC. Causal Evidence for the Role of Specific GABAergic Interneuron Types in Entorhinal Recruitment of Dentate Granule Cells. Scientific Reports. 2016 12; 6(1):36885. doi: 10.1038/srep36885.

Leung C, Cao F, Nguyen R, Joshi K, Aqrabawi AJ, Xia S, Cortez MA, Snead OC, Kim JC, Jia Z. Activation of Entorhinal Cortical Projections to the Dentate Gyrus Underlies Social Memory Retrieval. Cell Reports. 2018 5; 23(8):2379– 2391. doi: 10.1016/j.celrep.2018.04.073, publisher: Elsevier PMID: 29791849.

Leutgeb JK, Leutgeb S, Moser MB, Moser EI. Pattern Separation in the Dentate Gyrus and CA3 of the Hippocampus. Science. 2007 2; 315(5814):961–966. doi: 10.1126/science.1135801.

Liautard C, Scalmani P, Carriero G, de Curtis M, Franceschetti S, Mantegazza M. Hippocampal hyperexcitability and specific epileptiform activity in a mouse model of Dravet syndrome. Epilepsia. 2013 7; 54(7):1251–1261. doi: 10.1111/epi.12213.

Liu Y, Lopez-Santiago LF, Yuan Y, Jones JM, Zhang H, O’Malley HA, Patino GA, O’Brien JE, Rusconi R, Gupta A, Thompson RC, Natowicz MR, Meisler MH, Isom LL, Parent JM. Dravet syndrome patient-derived neurons suggest a novel epilepsy mechanism: Liu et al: DS Patient-Derived Neurons. Annals of Neurology. 2013 7; 74(1):128–139. doi: 10.1002/ana.23897.

Lu Y, Zhong C, Wang L, Wei P, He W, Huang K, Zhang Y, Zhan Y, Feng G, Wang L. Optogenetic dissection of ictal propagation in the hippocampal–entorhinal cortex structures. Nature Communications. 2016 4; 7(1):10962. doi: 10.1038/ncomms10962.

McHugh TJ, Jones MW, Quinn JJ, Balthasar N, Coppari R, Elmquist JK, Lowell BB, Fanselow MS, Wilson MA, Tonegawa S. Dentate Gyrus NMDA Receptors Mediate Rapid Pattern Separation in the Hippocampal Network. Science. 2007 7; 317(5834):94–99. doi: 10.1126/science.1140263.

Miller AR, Hawkins NA, McCollom CE, Kearney JA. Mapping genetic modifiers of survival in a mouse model of Dravet syndrome. Genes, brain, and behavior. 2014 2; 13(2):163–172. doi: 10.1111/gbb.12099, pMID: 24152123 PMCID: PMC3930200.

Mistry AM, Thompson CH, Miller AR, Vanoye CG, George AL, Kearney JA. Strain- and age-dependent hippocampal neuron sodium currents correlate with epilepsy severity in Dravet syndrome mice. Neurobiology of Disease. 2014 5; 65:1–11. doi: 10.1016/j.nbd.2014.01.006.

Neunuebel JP, Knierim JJ. Spatial Firing Correlates of Physiologically Distinct Cell Types of the Rat Dentate Gyrus. The Journal of Neuroscience. 2012 3; 32(11):3848–3858. doi: 10.1523/JNEUROSCI.6038-11.2012, pMID: 22423105 PMCID: PMC3321836.

Oakley JC, Kalume F, Yu FH, Scheuer T, Catterall WA. Temperature- and age-dependent seizures in a mouse model of severe myoclonic epilepsy in infancy. Proceedings of the National Academy of Sciences. 2009; 106(10):3994–3999. doi: 10.1073/pnas.0813330106.

Ogiwara I, Miyamoto H, Morita N, Atapour N, Mazaki E, Inoue I, Takeuchi T, Itohara S, Yanagawa Y, Obata K, Furuichi T, Hensch TK, Yamakawa K. Nav1.1 Localizes to Axons of Parvalbumin-Positive Inhibitory Interneurons: A Circuit Basis for Epileptic Seizures in Mice Carrying an Scn1a Gene Mutation. Journal of Neuroscience. 2007 5; 27(22):5903–5914. doi: 10.1523/JNEUROSCI.5270-06.2007.

Ogiwara I, Iwasato T, Miyamoto H, Iwata R, Yamagata T, Mazaki E, Yanagawa Y, Tamamaki N, Hensch TK, Itohara S, Yamakawa K. Nav1.1 haploinsufficiency in excitatory neurons ameliorates seizure-associated sudden death in a mouse model of Dravet syndrome. Human Molecular Genetics. 2013 12; 22(23):4784–4804. doi: 10.1093/hmg/ddt331, pMID: 23922229 PMCID: PMC3820136.

Osteen JD, Herzig V, Gilchrist J, Emrick JJ, Zhang C, Wang X, Castro J, Garcia-Caraballo S, Grundy L, Rychkov GY, Weyer AD, Dekan Z, Undheim EAB, Alewood P, Stucky CL, Brierley SM, Basbaum AI, Bosmans F, King GF, Julius D. Selective spider toxins reveal a role for Nav1.1 channel in mechanical pain. Nature. 2016 6; 534(7608):494–499. doi: 10.1038/nature17976, pMID: 27281198 PMCID: PMC4919188.

Pnevmatikakis EA. Analysis pipelines for calcium imaging data. Current Opinion in Neurobiology. 2019 4; 55:15–21. doi: 10.1016/j.conb.2018.11.004.

Pnevmatikakis EA, Soudry D, Gao Y, Machado TA, Merel J, Pfau D, Reardon T, Mu Y, Lacefield C, Yang W, Ahrens M, Bruno R, Jessell TM, Peterka DS, Yuste R, Paninski L. Simultaneous Denoising, Deconvolution, and Demixing of Calcium Imaging Data. Neuron. 2016 1; 89(2):285–299. doi: 10.1016/j.neuron.2015.11.037, pMID: 26774160 PMCID: PMC4881387.

Richards KL, Milligan CJ, Richardson RJ, Jancovski N, Grunnet M, Jacobson LH, Undheim EAB, Mobli M, Chow CY, Herzig V, Csoti A, Panyi G, Reid CA, King GF, Petrou S. Selective Na _V_ 1.1 activation rescues Dravet syndrome mice from seizures and premature death. Proceedings of the National Academy of Sciences. 2018 8; 115(34):E8077–E8085. doi: 10.1073/pnas.1804764115.

Rolls ET. A computational theory of episodic memory formation in the hippocampus. Behavioural Brain Research. 2010 12; 215(2):180–196. doi: 10.1016/j.bbr.2010.03.027.

Rubinstein M, Han S, Tai C, Westenbroek RE, Hunker A, Scheuer T, Catterall WA. Dissecting the phenotypes of Dravet syndrome by gene deletion. Brain. 2015 8; 138(8):2219–2233. doi: 10.1093/brain/awv142.

Salgueiro-Pereira AR, Duprat F, Pousinha PA, Loucif A, Douchamps V, Regondi C, Ayrault M, Eugie M, Stunault MI, Escayg A, Goutagny R, Gnatkovsky V, Frassoni C, Marie H, Bethus I, Mantegazza M. A two-hit story: Seizures and genetic mutation interaction sets phenotype severity in SCN1A epilepsies. Neurobiology of Disease. 2019 5; 125:31–44. doi: 10.1016/j.nbd.2019.01.006.

Senzai Y, Buzsáki G. Physiological properties and behavioral correlates of hippocampal granule cells and mossy cells. Neuron. 2017 2; 93(3):691–704.e5. doi: 10.1016/j.neuron.2016.12.011, pMID: 28132824 PMCID: PMC5293146.

Siegler Z, Barsi P, Neuwirth M, Jerney J, Kassay M, Janszky J, Paraicz E, Hegyi M, Fogarasi A. Hippocampal sclerosis in severe myoclonic epilepsy in infancy: a retrospective MRI study. Epilepsia. 2005 5; 46(5):704–708. doi: 10.1111/j.1528-1167.2005.41604.x, pMID: 15857436.

Stein RE, Kaplan JS, Li J, Catterall WA. Hippocampal deletion of NaV1.1 channels in mice causes thermal seizures and cognitive deficit characteristic of Dravet Syndrome. Proceedings of the National Academy of Sciences. 2019 8; 116(33):16571–16576. doi: 10.1073/pnas.1906833116, pMID: 31346088.

Stringer C, Pachitariu M, Steinmetz N, Reddy CB, Carandini M, Harris KD. Spontaneous behaviors drive multidimensional, brainwide activity. Science. 2019 4; 364(6437). https://science.sciencemag.org/content/364/6437/eaav7893, doi: 10.1126/science.aav7893, publisher: American Association for the Advancement of Science section: Research Article PMID: 31000656.

Sun Y, Paşca SP, Portmann T, Goold C, Worringer KA, Guan W, Chan KC, Gai H, Vogt D, Chen YJJ, Mao R, Chan K, Rubenstein JL, Madison DV, Hallmayer J, Froehlich-Santino WM, Bernstein JA, Dolmetsch RE. A deleterious Nav1.1 mutation selectively impairs telencephalic inhibitory neurons derived from Dravet Syndrome patients. eLife. 2016 7; 5:e13073. doi: 10.7554/eLife.13073.

Tai C, Abe Y, Westenbroek RE, Scheuer T, Catterall WA. Impaired excitability of somatostatin- and parvalbumin-expressing cortical interneurons in a mouse model of Dravet syndrome. Proceedings of the National Academy of Sciences. 2014 7; 111(30):E3139–E3148. doi: 10.1073/pnas.1411131111.

Tiefes AM, Hartlieb T, Tacke M, von Stülpnagel-Steinbeis C, Larsen LHG, Hao Q, Dahl HA, Neubauer BA, Gerstl L, Kudernatsch M, Kluger GJ, Borggraefe I. Mesial Temporal Sclerosis in SCN1A-Related Epilepsy: Two Long-Term EEG Case Studies. Clinical EEG and neuroscience. 2019 7; 50(4):267–272. doi: 10.1177/1550059418794347, pMID: 30117335.

Tran CH, Vaiana M, Nakuci J, Somarowthu A, Goff KM, Goldstein N, Murthy P, Muldoon SF, Goldberg EM. Interneuron Desynchronization Precedes Seizures in a Mouse Model of Dravet Syndrome. Journal of Neuroscience. 2020 3; 40(13):2764–2775. doi: 10.1523/JNEUROSCI.2370-19.2020, publisher: Society for Neuroscience section: Research Articles PMID: 32102923.

Treves A, Tashiro A, Witter MP, Moser EI. What is the mammalian dentate gyrus good for? Neuroscience. 2008 7; 154(4):1155–1172. doi: 10.1016/j.neuroscience.2008.04.073.

Tsai MS, Lee ML, Chang CY, Fan HH, Yu IS, Chen YT, You JY, Chen CY, Chang FC, Hsiao JH, Khorkova O, Liou HH, Yanagawa Y, Lee LJ, Lin SW. Functional and structural deficits of the dentate gyrus network coincide with emerging spontaneous seizures in an Scn1a mutant Dravet Syndrome model during development. Neurobiology of Disease. 2015 5; 77:35–48. doi: 10.1016/j.nbd.2015.02.010.

Van Poppel K, Patay Z, Roberts D, Clarke DF, McGregor A, Perkins FF, Wheless JW. Mesial Temporal Sclerosis in a Cohort of Children With SCN1A Gene Mutation. Journal of Child Neurology. 2012 7; 27(7):893–897. doi: 10.1177/0883073811435325.

Xiong G, Metheny H, Johnson BN, Cohen AS. A Comparison of Different Slicing Planes in Preservation of Major Hippocampal Pathway Fibers in the Mouse. Frontiers in Neuroanatomy. 2017 11; 11. https://www.ncbi.nlm.nih.gov/pmc/articles/PMC5696601/, doi: 10.3389/fnana.2017.00107, pMID: 29201002 PMCID: PMC5696601.

Xu JH, Tang FR. Voltage-Dependent Calcium Channels, Calcium Binding Proteins, and Their Interaction in the Pathological Process of Epilepsy. International Journal of Molecular Sciences. 2018 9; 19(9). https://www.ncbi.nlm.nih.gov/pmc/articles/PMC6164075/, doi: 10.3390/ijms19092735, pMID: 30213136 PMCID: PMC6164075.

Yamagata T, Raveau M, Kobayashi K, Miyamoto H, Tatsukawa T, Ogiwara I, Itohara S, Hensch TK, Yamakawa K. CRISPR/dCas9-based Scn1a gene activation in inhibitory neurons ameliorates epileptic and behavioral phenotypes of Dravet syndrome model mice. Neurobiology of Disease. 2020 7; 141:104954. doi: 10.1016/j.nbd.2020.104954.

Yu EP, Dengler CG, Frausto SF, Putt ME, Yue C, Takano H, Coulter DA. Protracted Postnatal Development of Sparse, Specific Dentate Granule Cell Activation in the Mouse Hippocampus. The Journal of Neuroscience. 2013 2; 33(7):2947–2960. doi: 10.1523/JNEUROSCI.1868-12.2013, pMID: 23407953 PMCID: PMC3711669.

Yu FH, Mantegazza M, Westenbroek RE, Robbins CA, Kalume F, Burton KA, Spain WJ, McKnight GS, Scheuer T, Catterall WA. Reduced sodium current in GABAergic interneurons in a mouse model of severe myoclonic epilepsy in infancy. Nature Neuroscience. 2006 9; 9(9):1142–1149. doi: 10.1038/nn1754.

